# A Clock and Wavefront Self-Organizing Model Recreates the Dynamics of Mouse Somitogenesis in-vivo and in-vitro

**DOI:** 10.1101/2023.01.18.524516

**Authors:** Julie Klepstad, Luciano Marcon

## Abstract

During mouse development, presomitic mesoderm cells synchronize Wnt and Notch oscillations, creating sequential phase waves that pattern somites. Traditional somitogenesis models attribute phase waves to global signals that control the frequency of oscillations. However, increasing evidence suggests that they could arise in a self-organizing manner. Here, we introduce the Sevilletor, a novel reaction-diffusion system that serves as a framework to compare different somitogenesis patterning hypotheses. Using this framework, we propose the Clock and Wavefront Self-Organizing model, the first somitogenesis model that generates phase waves via local cell to cell communication independent of global frequency gradients. The model recapitulates the change in relative phase of Wnt and Notch observed during mouse somitogenesis and the formation of multiple phase waves observed upon ectopic expansion of posterior signals. Moreover, it provides a theoretical basis for understanding the excitability of mouse presomitic mesoderm cells observed in vitro.

## Introduction

During embryonic development, precise coordination of cell behaviors in both time and space is essential to generate robust gene expression patterns at the tissue level. A remarkable example of this coordination can be observed during somitogenesis, the formation of body segment precursors. In vertebrates, this process is characterized by waves of gene expression that propagate from the posterior tip of the tail to the anterior side, resulting in the sequential formation of somites (Figure 1A). These waves are generated by synchronizing oscillations driven by the segmentation clock, a genetic network of presomitic mesoderm (PSM) cells that pattern somites in a rhythmic manner. Previous studies have demonstrated that the core of the segmentation clock is implemented by delayed negative feedbacks [26] and that the waves arise from a phase shift of the clock along the anterior-posterior axis [28, 3]. The specific mechanism responsible for the spatial synchronization of oscillations, however, remains a subject of debate. One possibility is that the oscillations are modulated by global coordinating signals, such as anterior-posterior frequency profiles. Another possibility is that the oscillations are synchronized through local cell communication.

**Figure 1:**
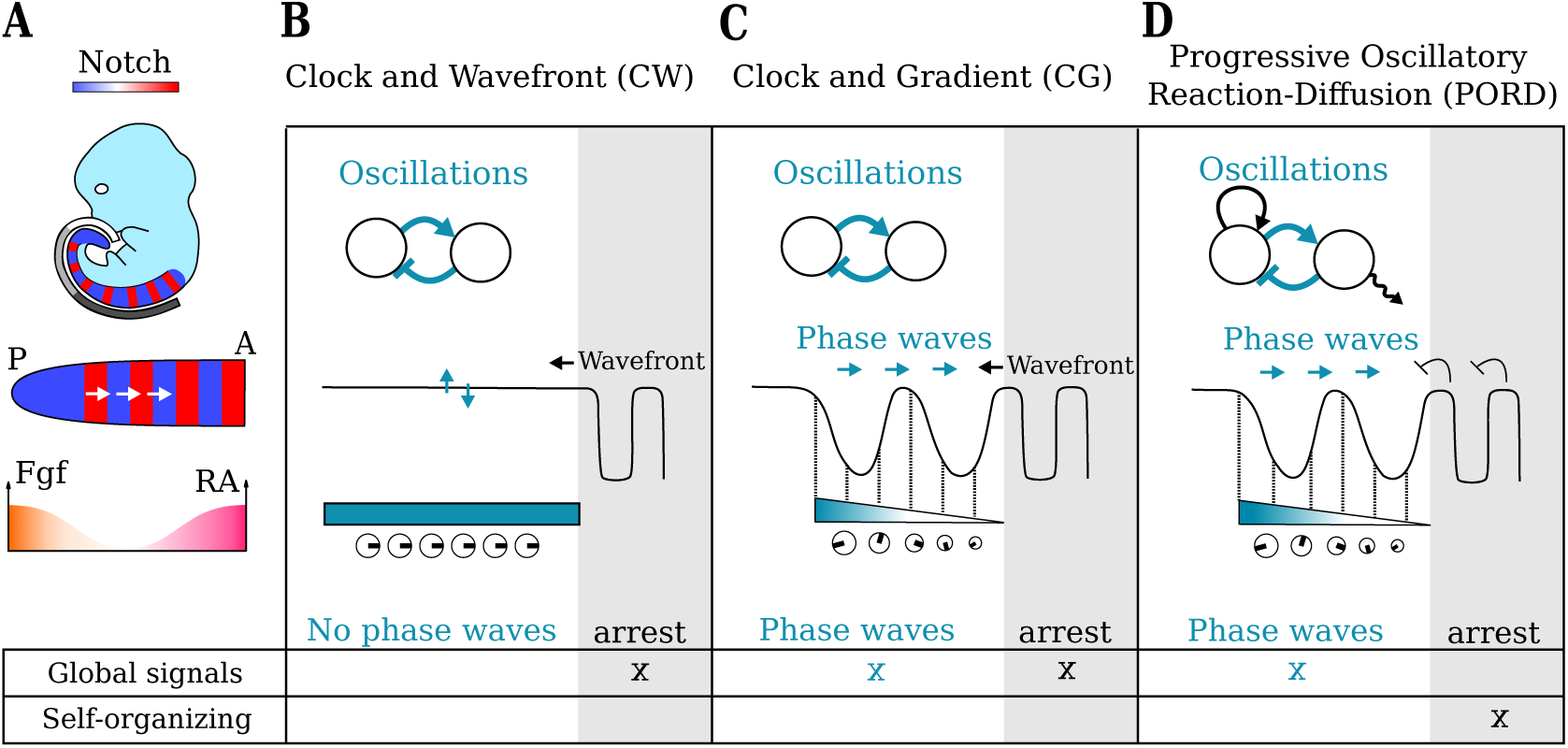
Introductory Figure. **A)** During somitogenesis, coordinated genetic oscillation gives rise to phase waves traversing the embryo from the posterior (P) to the anterior (A) side. This process is orchestrated by anterior-posterior signals such as Fgf and RA. **B-D)** In previous somitogenesis models, oscillations can emerge from a delayed negative feedback (blue arrows) with a phase (black line in 1D graph) that is promoted by different spatial frequency profiles (blue/white regions and clocks). On the bottom: a table indicates whether the formation and arrest of phase waves is controlled by global signals or is a self-organizing process. **B)** In the simple Clock and Wavefront model (CW) [8], all the cells have the same frequency (blue rectangle and identical clocks) giving rise to homogeneous phase oscillations. Global signals define a moving determination front that arrests the oscillations generating a periodic somite pattern. **C)** In the Clock and Gradient model (CG) [31, 17], in addition, a monotonically decreasing frequency profile (blue gradient and clocks with varying sizes) gives rise to phase waves. **D)** In the Progressive Oscillatory Reaction-Diffusion Model (PORD) [9], phase waves are formed by a graded frequency profile but the arrest of oscillations is a self-organizing process.

The Clock and Wavefront (CW) model [8] was the first theoretical study to suggest that oscillation in the posterior part of the PSM (clock) controls somitogenesis. According to this model, when cells exit this posterior region they enter a determination front (wavefront) defined by global positional information signals [53] undergoing rapid changes governed by the phase of the clock to form periodic somite patterns (Figure 1B). The main ideas of the CW model have received strong experimental support in Chick, Zebrafish, and Mouse, where it has been shown that the posterior PSM exhibits homogeneous oscillations of Notch, Wnt and Fgf signaling [1, 36, 38]. Moreover, experimental evidence shows that posterior signaling gradients of Wnt and Fgf allow cells to oscillate at the posterior tip of the tail [10, 33, 1], while Retinoic acid (RA) forms an anterior gradient that localizes within newly formed somites and promotes differentiation [11, 1, 36], see Figure 1A.

In its original formulation, however, the CW model fails to account for the formation of phase waves observed during vertebrate somitogenesis. A popular reincarnation of the CW model that addresses this issue is the Clock and Gradient model (CG), which assumes that the frequency of the clock slows down gradually from posterior to anterior [31, 17]. This decreasing frequency profile promotes a spatial alternation of phases, thereby giving rise to waves (Figure 1C). Adding local coupling to the CG model enhances robustness and scales the overall frequency [31, 17]. However, it is important to note that the formation of wave patterns in these models is tightly linked to the slope of the frequency profile. For instance, it has been proposed that slightly shallower profiles in Zebrafish can promote the formation of multiple phase waves in comparison with mouse [37]. In essence, in the CG model, both the arrest of oscillations and the formation of phase waves are under the control of global signals (Figure 1C). Intriguingly, genetic manipulations in mouse have revealed that multiple phase waves can be formed upon the ectopic activation of Wnt signaling over the whole PSM, which ensues saturation of posterior signaling and flattening of the gradients [3]. This observation suggests that the formation of phase waves may not be explicitly governed by monotonically decreasing gradients as in the CG model, but could display a degree of self-organization.

The idea that PSM cells can coordinate oscillations in a self-organizing manner has been explored in the Progressive Oscillatory Reaction-Diffusion (PORD) model [9]. In contrast to the CW and CG models, where new somites formed sequentially due to the progressive movement of the wavefront, in the PORD model, new somites form in a self-organizing manner through a relay mechanism triggered by the last formed somite (Figure 1D). Thus, somite formation becomes independent of posterior signaling gradients and instead relies on a pre-patterned somite. When a modulation promoted by a posterior morphogen gradient is added to the model, it can also recapitulate the formation of phase waves in a manner similar to the CG model [9]. In summary, in the PORD model, the arrest of oscillations is driven by a self-organizing relay mechanism while the formation of phase waves is still under the control of global signals (Figure 1D)

Therefore, thus far, none of the existing somitogenesis models has explored the possibility that phase waves form in a self-organizing manner independent of global signals. Nevertheless, recent experimental studies have challenged this notion by culturing two-dimensional tail explants in vitro [25, 49, 18]. These explants can generate circular phase waves that resemble somitogenesis despite lack of embryonic signals and upon significant cellular rearrangements. The presence [18] or absence [25] of gradients that could potentially direct the formation of these circular waves remains unclear. However, wave formation has also been observed in more heterogeneous cultures obtained by mixing cells from different tailbud explants [18]. This highlights that PSM cells possess the self-organizing capacity to synchronize oscillations and form phase waves. How somitogenesis models can capture this self-organizing behavior remains to be explored.

The behavior of isolated PSM cells in vitro has also provided insights into the role played by cell communication during Zebrafish and mouse somitogenesis. In Zebrafish, isolated cells exhibit default oscillatory behavior and local cell communication is necessary to synchronize oscillations [27, 28, 52, 36], as in classical Kuramoto models [23]. In mouse, isolated cells display a default stable behavior and can be induced to oscillate by increasing cell density [18]. This shows that, in mouse, local cell communication is essential for both initiating oscillations at the individual cell level and synchronizing oscillations at the tissue level [34]; suggesting that excitatory behaviors could play a significant role in phase wave formation [18]. Nevertheless, it is still unclear whether these differences depend on specific in-vitro culture conditions. Indeed, a recent study has shown that the oscillations of isolated Zebrafish cells cease after a certain number of cycles, which varies based on the anterior-posterior position [42]. On the other hand, isolated mouse PSM cells can be induced to oscillate also when embedded in a BSA substrate or by inhibiting actin polymerization with latrunculin A [18].

Here, we present a novel minimal theoretical framework to explore the role of global coordinating signals and self-organization in mouse somitogenesis. Our framework consists of two main components: a core reaction-diffusion system implemented by a delayed negative feedback between two self-enhancing genes, and a posterior signaling gradient. At the single cell level, this reaction-diffusion system can produce oscillations or bi-stability. However, when local cell-to-cell coupling occurs via diffusion, various self-organizing behaviors can emerge at the tissue level, including lateral inhibition, rotating waves, homogeneous patterns, and bi-stable patterns. While similar patterns have been observed in previous reaction-diffusion models [43], one of the novelties of our system lies in its ability to generate this wide range of self-organizing behaviors by adjusting a single reaction parameter. Additionally, our system exhibits a previously undescribed patterning behavior: the formation of periodic phase waves through the excitation of a bistable state. These periodic phase waves include spirals and resemble the patterns formed by models of the Belousov–Zhabotinsky chemical reaction [54], such as the Brusselator [41, 35] and the Oregonator [13], which were developed in Brussels (Belgium) and Oregon (USA) respectively. Continuing the tradition, we named our theoretical framework the Sevilletor, as it was developed in Seville (Spain).

We demonstrate that the Sevilletor framework can recapitulate the principal theoretical models proposed to explain somitogenesis with minimal parameter changes. In addition, we present a novel somitogenesis model that for the first time explains phase wave formation as a guided self-organizing process. We name this model the Clock and Wavefront Self-Organizing model (CWS), since it extends the CW model with an excitable self-organizing region. Notably, the addition of this region can also explain the change in the relative phase between Notch and Wnt observed in the middle part of the mouse tail [2, 46]. By performing two-dimensional simulations of the developing tail, we demonstrate that the CWS can form multiple phase waves upon the ectopic expansion of posterior signals as observed in mutants [3]. Lastly, we employ our theoretical framework to investigate the behavior of PSM cells within virtual explants in each model. Our analysis reveals that only CWS explants can generate phase waves in the absence of gradients and the inherent phase values present in the tail. Overall, we demonstrate that the CWS model is the first somitogenesis hypothesis that can explain the self-organizing potential of the PSM both in-vivo and in-vitro.

## Results

We begin the results section providing a summary of a detailed theoretical analysis presented in the method section, where we use complex systems theory and numerical simulations to introduce and characterize a new reaction-diffusion system called Sevilletor. The rest of the result section focuses on the application of this system to study somitogenesis. Specifically, we use these equations to investigate the qualitative behavior of previous somitogenesis models and to propose a novel somitogenesis model where phase waves are formed in a self-organized manner.

### Summary of the theoretical analysis of the Sevilletor system

We devised a minimal equation system called Sevilletor that couples two self-enhancing reactants *u* and *v* with a negative feedback, and limits the deviation of concentrations with cubic saturation terms, see Figure 2A and equations (1) and (2), introduced in detail in the method section. On the one hand, the negative feedback between the two reactants (*k*_3_ and *k*_4_) in the model promotes sustained oscillations like the one generated by single reactant models with a delayed negative feedback. On the other hand, varying the relative self-enhancement strength of *u* and *v* (*k*_1_ and *k*_2_) promotes a bifurcation from an oscillatory state around an unstable point to a bistable state (Figure 2A-C), adding four additional steady states (Figure 2F).

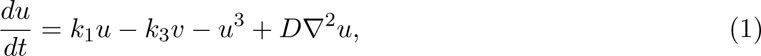

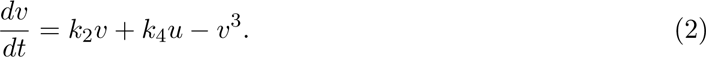

**Figure 2:**
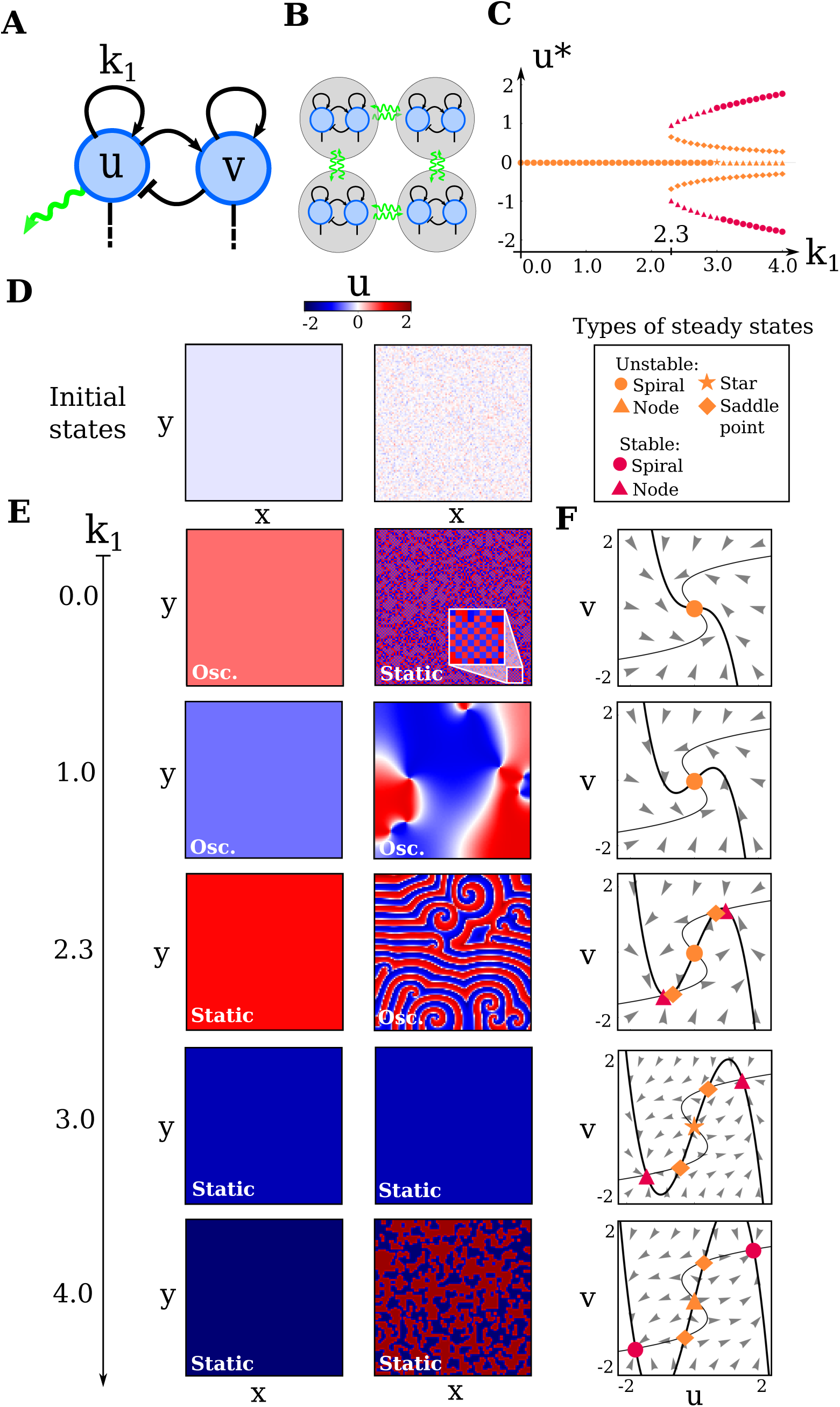
Patterning behavior and bifurcations in the Sevilletor equations. **A)** The Sevilletor equations couple two self-enhancing reactants *u* and *v* with a negative feedback. *k*_1_ defines the strength of the self-enhancement of *u*, while *k*_2_,_3_,_4_ = 1. **B)** Diffusion of *u* couples the behavior of four neighboring cells (green arrows). **C)** Bifurcation diagram of the Sevilletor equations for *k*_1_. **D)** Initial conditions of *u* used for the simulations in (E): homogeneous initial concentrations on the left and random initial conditions on the right. **E)** From top to bottom, two dimensional simulations with increasing values of *k*_1_. Left column, the model produced homogeneous oscillatory patterns for 0 *≤ k*_1_ *<* 2.3 and static homogeneous patterns for *k*_1_ *≥* 2.3 with homogeneous initial concentrations. Right column, starting from random initial conditions the model forms lateral inhibition patterns with *k*_1_ = 0.0 (the zoomed region shows a chessboard pattern), rotating phase waves with *k*_1_ = 1.0, periodic waves with spirals *k*_1_ = 2.3, homogeneous patterns for *k*_1_ = 3.0 and bi-stable salt and pepper pattern for *k*_1_ = 4.0 (Movie 1). **F)** From top to bottom, phase spaces with increasing values of *k*_1_ = [0.0, 1.0, 2.3, 3.0, 4.0]. The thick line shows the nullcline of *u* and thin line the nullcline of *v*, markers show the steady states (see legend above) and arrowheads the direction of the vectorfield. The shape of the nullcline of *u* (thick line) changes as *k*_1_ increases, undergoing a bifurcation at *k*_1_ = 2.3 gaining two stable states (red markers) near two unstable states (orange markers).

In addition, we observe that for different self-enhancement strengths, diffusion of *u* between cells can give rise to five different patterning behaviors: lateral inhibition patterns, rotating wave patterns, periodic wave patterns with spiral formation, homogeneous patterns and bi-stable frozen patterns (right column in Figure 2E, Figure 3H-J and Movie 1). An analysis with two-cell simulations revealed that diffusion plays a different role in each of these scenarios, as it can be seen by comparing the trajectories with and without diffusion in phase space (Figure 3B-F and Movies 2 and 3). It can be either stabilizing (lateral inhibition) to freeze the oscillations of neighboring cells in opposite phase generating chessboard patterns, synchronizing to generate rotating waves, or destabilizing. In the latter case cells are in a default bistable state and diffusion excite cells to oscillate when they are in opposite phase, generating periodic phase waves, Figure 3D and Movie 3. This excitable self-organizing patterning behavior has not been described previously and differs from classical diffusion-driven Turing instability [50] or periodic wave patterns observed in Kuramoto models [6, 51].

**Figure 3:**
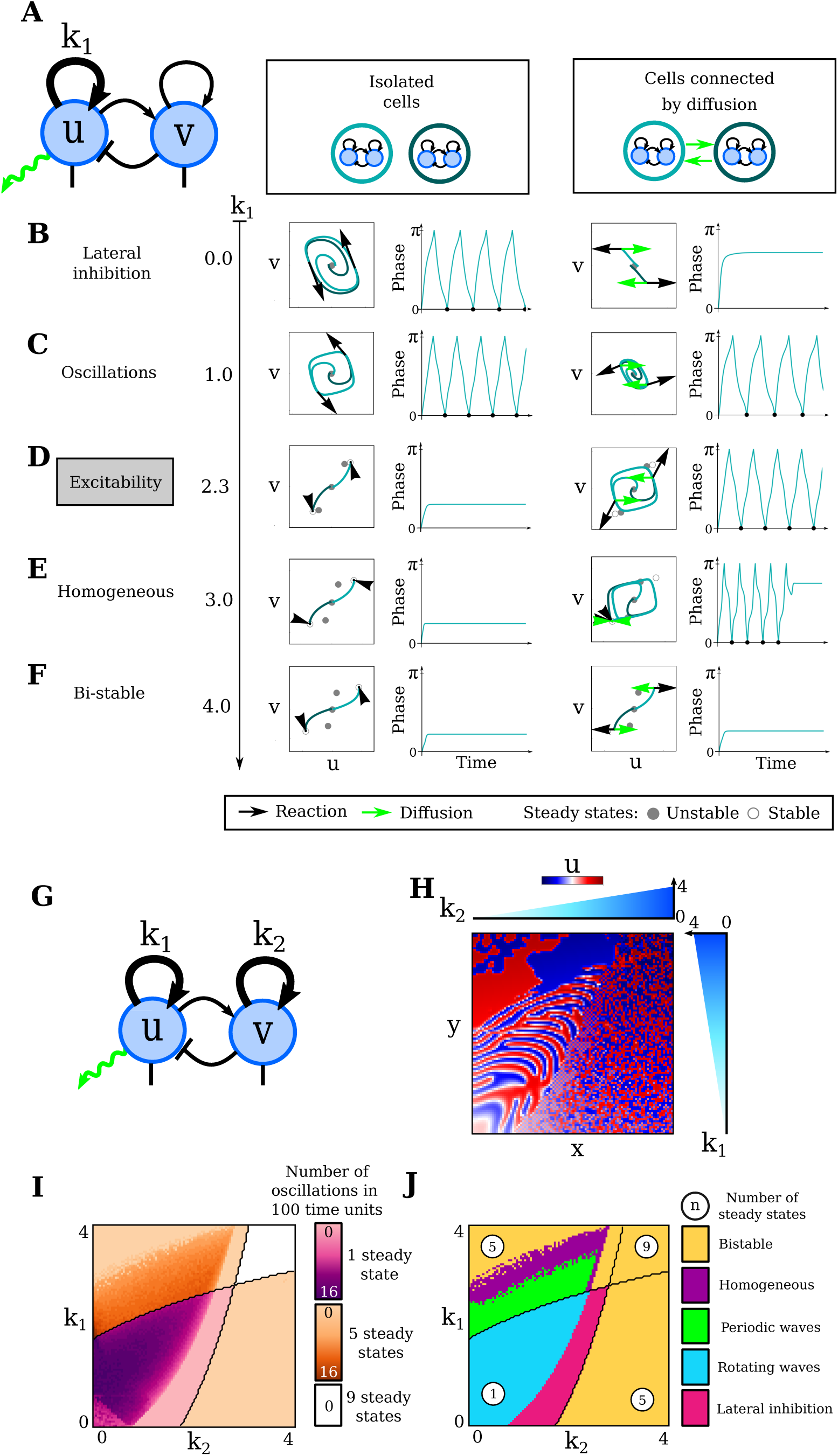
The effect of variables *k*_1_, *k*_2_ and diffusion in the Sevilletor equations. **A-F)** Varying *k*_1_ promotes different dynamical behaviors in two cell simulations, where the cells have opposite phases, with and without diffusion. Initial conditions are (*u, v*) = (*−*0.1, 0) in the first cell (dark teal color) and (*u, v*) = (0.1, 0) in the second cell (light teal color). Left graphs show the trajectory of the two cells, graph on the right the phase of one cell over time. Black and green arrows highlight the contribution of reaction and diffusion respectively. **B)** With *k*_1_ = 0 isolated cells undergo sustained oscillations but stop oscillating when coupled by diffusion, because reaction (black arrows) is counterbalanced by diffusion (green arrow) (Movie 2). **C)** With *k*_1_ = 1 isolated cells undergo sustained oscillations and continue oscillating when coupled by diffusion. **D)** With *k*_1_ = 2.3 isolated cells are frozen in a bi-stable regime (black arrows pointing to stable states) but can excite each other to oscillate when coupled by diffusion (green arrow), which pushes each each cell towards the trajectory of the nearest unstable point (gray dots) generating a new limit cycle (Movie 3). **E)** With *k*_1_ = 3 isolated cells are frozen in a bi-stable regime but can excite each other to oscillate when coupled by diffusion (green arrow) synchronizing the oscillations and eventually stopping. **F)** With *k*_1_ = 4 both isolated and coupled cells are frozen in a bi-stable regime. **G-J)** Bifurcation diagrams for *k*_1_ and *k*_2_ and associated numerical simulations. **H)** Two-dimensional simulation with graded values of *k*_1_ along the y-axis and *k*_2_ along the x-axis that recapitulates all patterning behaviors of the Sevilletor model (Movie 5). **I-J)** The number of oscillations quantified in two cell simulations for different values of *k*_1_ and *k*_2_ (color intensity of each point in I), extends the bifurcation diagram for *k*_1_ and *k*_2_ showing the different dynamical patterning regimes promoted by diffusion in the Sevilletor model (colored regions in J). Black lines in I and J show bifurcations in the model without diffusion.

In the following section, we show that these different dynamical behaviors are amenable to simulate the main patterning hypotheses proposed to study mouse somitogenesis.

### Sevilletor as a framework to study somitogenesis

Our goal is to compare the qualitative patterning behavior of different somitogenesis models by exploiting the dynamical patterning regimes exhibited by the Sevilletor equations. Our focus is solely to explain the patterning dynamics observed during mouse somitogenesis. A deeper analysis of the differences between the mouse and Zebrafish presomitic mesoderm is beyond the scope of this study but it is discussed for future work in the discussion section.

Within this context, our objective is to capture the qualitative aspects that control the emergence of oscillations, phase waves, and the arrest of oscillations in different models, rather than replicating the quantitative details of each scenario. To do so, we modify the Sevilletor equations by adding two spatial functions R (*regions*) and PG (*posterior gradient*) that promote distinct modulations along the anterior posterior axis of the developing tail. We rewrite the equations (1) and (2) as:

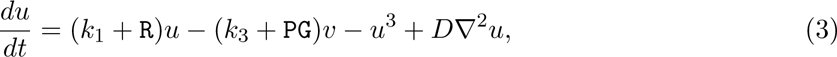

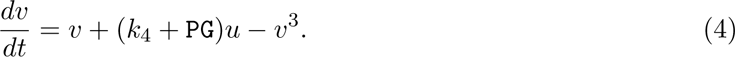

We simulate these equations on a 2D growing rectangular grid of virtual cells that elongates with constant speed along the y-axis, which represents the anterior-posterior axis of the developing tail (details in the methods section). In the model, tail growth is implemented by proliferation of the posterior-most cells, which generate a new line of cells that inherit the concentrations of the two reactants *u* and *v*. The functions R and PG have sigmoidal spatial profiles along the y axis and are defined as 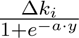, where *y* represents an anterior-posterior spatial coordinate, Δ*k_i_* the amplitude of the sigmoid function and *a* its steepness.

The spatial function R is used to regionalize the anterior-posterior axis into discrete regions with different dynamical regimes. This is done with a steep sigmoid function obtained with *a* = 1 that modulates the parameter *k*_1_ in a step-like fashion in space. Indeed, the theoretical analysis presented in Figure 2 and Figure 3 show that *k*_1_ is a key parameter that promotes bifurcations and changes in dynamical system behavior. This approach allows us, for example, to promote a bifurcation from a posterior oscillatory regime (*k*_1_ *<* 2.3) to an anterior bi-stable regime (*k*_1_ *>* 2.3) providing the basis to investigate how cells become committed to form somites as they move anteriorly (Figure 4A-B). This modulation is sufficient to devise a basic dynamical model that captures the features of the Clock and Wavefront model (CW) [8] (Figure 4C). This can be done by setting *k*_1_ = 4 and 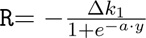, where *y* is the anterior-posterior coordinate from the posterior boundary, Δ*k*_1_ = 3 and *a* = 1, defining a moving wavefront that promotes the commitment of cells to a specific phase. This approach is similar to the bifurcations presented in [15, 20] and in models of neural tube patterning [39].

**Figure 4:**
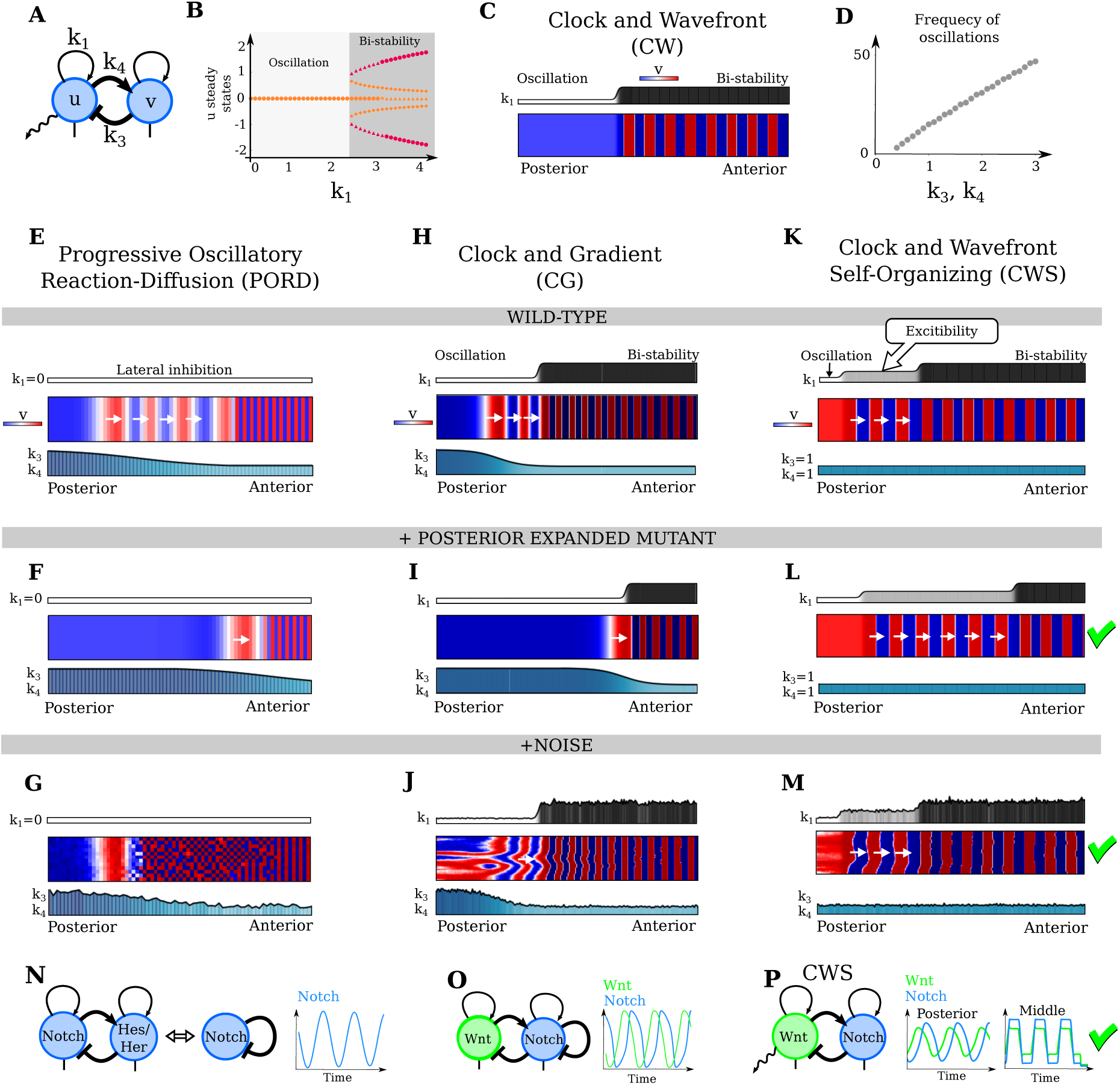
Classical somitogenesis patterning hypotheses and the Clock and Wavefront Self-Organizing model in the Sevilletor framework. **A-B)** Varying the value of *k*_1_ changes from an oscillatory to a bistable state as shown by the bifurcation diagram (B). **C)** The Clock and Wavefront model (CW) consider a bifurcation from oscillatory regime in the posterior side (*k*_1_ = 1) to a bistable anterior side (*k*_1_ = 4) that is defined by a wavefront. As the tail grows, cells enter the wavefront giving rise to a periodic bi-stable somite pattern. **D)** The frequency of oscillation for *k*_1_ *<* 2.3 increases linearly with the strength of the negative feedback loop (*k*_3_ and *k*_4_). **E-M)** Two-dimensional simulations of the PORD, CG and CWS somitogenesis models in the Sevilletor framework. **E,H,K)** Simulated patterns of *v* in each model promoted by different dynamical regimes *k*_1_ (white-black) and frequency gradients promoted by *k*_3_ and *k*_4_ (white-blue). White arrows show phase waves. **F,I,L)** Simulated patterns of *v* in an expanded posterior mutant in each model. **G,J,M)** Simulated patterns of *v* with 5% noise added to concentrations and 0.5% to parameters. **E-G)** A unique dynamical regime with *k*_1_ = 0 implements the PORD model generating thin stripes sequentially via a relay mechanism triggered by a pre-patterned anterior somite (Movie 6). A frequency gradient (blue gradient of *k*_3_ and *k*_4_) directs phase wave formation. Less phase waves are formed in the expanded posterior mutant and the somite pattern is fragile to noise. **H-J)** The Clock and Gradient model (CG) adds an anterior-posterior gradient to the CW model (blue gradient of *k*_3_ and *k*_4_) promoting a frequency profile that drives phase wave formation (Movie 7). Less phase waves are formed in the expanded posterior mutant and noise in the frequency profile disrupts the pattern over time. **K-M)** The Clock and Wavefront Self-Organizing model (CWS) adds a middle section with an excitability regime to the CW model (excitable gray region with *k*_1_ = 2.3) where phase waves are formed without a frequency gradient (same *k*_3_ and *k*_4_ value in blue), see Movie 8. More phase waves are formed in the expanded posterior mutant as seen in [3] and the model has a general good robustness to noise. **N)** The negative feedback between *u* and *v* can be interpreted as a negative feedback between Notch and Hes/Her, equivalent to a delayed inhibition of Notch. **O)** The delayed inhibition of Notch can coexist with a feedback between Notch and Wnt producing oscillations of Wnt and Notch that are coupled. **P)** The simple CWS model considers only the feedback between Wnt and Notch, which is sufficient to drive oscillations. On the right, average Wnt (*u* lime) and Notch (*v* cyan) values in the posterior and middle part of the CWS model (K) showing a relative phase change from out of phase to in phase oscillation as observed in experiments [2, 46].

The spatial function PG in equations (3) and (4) represents the effect of posteriorly graded signals and it is defined as 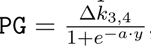, where *y* is the anterior-posterior coordinate from the posterior boundary and *a* = 0.1 to promote a graded spatial modulation with low steepness. The function controls the strength of the negative feedback between *u* and *v*, which similarly to the strength of the delayed negative feedback in single reactant models [31, 17] controls the frequency of oscillations. Specifically, the parameter Δ*k*_3_,_4_ promotes a change in *k*_3_ and *k*_4_, whose strengths are linearly correlated with oscillation frequency (Figure 4D).

In the next section, we demonstrate that modulations promoted by R and PG are sufficient to capture some of the main features of previous somitogenesis models. In addition, we show that they can be used to devise a novel somitogenesis model that extends the Clock and Wavefront model with an intermediate region where presomitic mesoderm cells are in a self-organizing excitable regime.

### A Sevilletor implementation of the PORD model

The first somitogenesis patterning hypothesis we explore is the Progressive Oscillatory Reaction-Diffusion (PORD) model from [9] (supplementary Section S6). This model generates sequential somite patterns through a relay self-organizing mechanism driven by diffusion. To form somites, the PORD model requires initial conditions with an anterior line of cells in the same high phase, triggering the relay mechanism throughout the tail. Subsequent studies reformulated the PORD model, triggering relay mechanisms with a posterior gradient without the need for a pre-patterned somite [40, 24].

All PORD models share the common characteristic of generating periodic somite patterns where the peaks of at least one reactant span over few cells. Analyzing the original PORD model from [9] through two-cell simulations (similar to Figure 3B), we discovered that these peaks are formed by freezing oscillations of neighboring cells in opposite phase (Figure S8) like in the lateral inhibition case of the Sevilletor equations. Therefore, to recreate the PORD model within the Sevilletor equations, we assume all cells in the tail are in an oscillatory regime without regionalization (R=0), with parameters *k*_1_ = 0 and *D* = 1 to trigger lateral inhibition (Figure S8B). Starting from an anterior pre-patterned somite, the model produces somites with a one-cell width sequentially due to a relay mechanism (Movie 2) similar to the original PORD model (Figure 4E). Introducing a graded frequency modulation with the function PG with Δ*k*_3_,_4_ = 0.3 and *a* = 0.1 generates phase waves as in the original PORD model (Figure 4E and Movie 6).

Simulations predict that expanding the posterior gradient leads to phase wave loss due to the posterior flattening of the frequency profile and a steeper drop on the anterior side (Figure 4F). This prediction contrasts with observed phenotypes in mice with a beta-catenin gain-of-function allele [3], where multiple phase waves form despite overall upregulation of Wnt and Fgf signaling. According to authors of [3], no appreciable posterior-to-anterior gradients can be detected, as revealed by beta-catenin immuno stainings and expression of direct Wnt and Fgf signaling targets.

Lastly, our PORD model implementation confirms its fragility to spatial noise, consistent with [40]. In both the original PORD model and our implementation, noise disrupts the periodic somite patterns resulting in a salt-and-pepper pattern (Figure 4G and Figure S9). In summary, while the lateral inhibition regime of the PORD model explains anterior oscillation arrest through local cell communication without changing the oscillatory regime, it forms periodic patterns with peaks spanning few cells that are sensitive to noise.

### A Sevilletor implementation of the Clock and Gradient model

The second somitogenesis patterning hypothesis that we explore is the Clock and Gradient (CG) model, a popular reincarnation of the Clock and Wavefront (CW) model [8] where the frequency of the segmentation clock is modulated along the anterior-posterior axis promoting phase wave formation. Typically, the CG model has been represented as a series of phase oscillators [31, 17] with frequencies of oscillations determined by a spatial frequency profile decreasing progressively anteriorly [31]. This model also incorporated local coupling between oscillators that promoted higher collective frequency and robustness to noise [31], though it did not contribute to phase wave formation, which was determined primarily by the frequency profile slope [37]. Finally, the arrest of oscillations was mainly represented with an anterior frequency approaching zero, interpreted as a phenomenological mechanistic transition to arrest oscillations driven by a wavefront [31] or by a transition promoted by two opposing signals [21].

In our Sevilletor version of the CG model, both the initiation and arrest of oscillations are emergent features of the dynamical system (Figure 4H, Movie 7). A spatial frequency profile that induces phase waves is introduced by decreasing the feedback strength between *u* and *v* with the posteriorly graded function PG (Δ*k*_3_,_4_ = 2, *a* = 0.1) similarly to the PORD model.

The oscillatory regime in the posterior tail for *k*_1_ = 1 is an emergent phenomenon driven by the negative feedback between *u* and *v*, leading to sustained cell-autonomous oscillations, equivalent to phase oscillators [31, 17, 21] or those in delayed negative feedback models [26]. If cells are coupled along the anterior-posterior axis by diffusion with *D* = 0.3, they exhibit robust collective oscillations resistant to noise. If started from random phases, the coupling of oscillations would cause rotating phase wave patterns to be formed, similar to pinwheel waves in Kuramoto models [6] as in the CG model in [51].

Consistent with previous CG model implementations, the frequency profile generates phase waves that become thinner moving anteriorly and freeze into periodic patterns entering the bistable determination front, marking a bifurcation between an oscillatory regime and bistability. This is obtained with a bifurcation promoted by R with Δ*k*_1_ = 3 and *a* = 1, using a similar approach as other body segmentation models [15, 20].

The phase waves form in a purely cell-autonomous manner even for *D* = 0, but with diffusion (*D >* 0) the wave formation is more robust to noise applied to the phases, as showed in [17]. However, applying noise to the posterior gradient PG causes somite pattern disorganization over time (Figure 4J and Figure S10). Finally, the model predicts a loss of phase waves in mutants with posterior signal expansion and flattening, contrary to experiments [3] (Figure 4I).

### The Clock and Wavefront Self-Organizing model

Motivated by the observation that in both the PORD and CG models, phase waves are generated by a global frequency profile whose slope controls the number of phase waves formed, and precision correlates with patterning robustness, we investigated whether the Clock and Wavefront model can be extended to produce phase waves via local cell-to-cell communication.

The foundational concept of this new model aligns with the basis of the Clock and Wavefront (Figure 4C) and the Clock and Gradient models, where cells oscillate posteriorly, transitioning to bistability through a determination wavefront. In our novel Clock and Wavefront Self-Organizing (CWS) model (Figure 4K), however, we introduce a third discrete region defined by the regionalizing function R, where cells have intermediate values of *k*_1_. In the basic formulation of the CWS model, we omit graded signals (PG) to emphasize the model’s capacity of forming phase waves independently of graded modulations. Nevertheless, in the next section, we demonstrate that the model can be adjusted to incorporate a graded modulation capturing the changes in phase wave width observed in vivo.

A key motivation for the CWS model is to devise a model that can generate phase waves independently of specific shapes of posterior gradients [3] and can recapitulate the excitability of mouse PSM cells observed in vitro [18]. The CWS model is implemented by regionalizing the tail in three parts with two steep changes in the parameter *k*_1_ promoted by a function R from *k*_1_ = 2.3. The function R that regionalizes the tail in three discrete regions is defined with a posterior and an anterior sigmoid function with steep changes defined as:

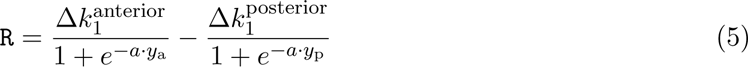

where *y*_p_ and *y*_a_ are the *y* coordinate from the posterior and anterior boundary respectively, Δ*k*_1_,_anterior_ = 1.7 to promote bistability, Δ*k*_1_,_posterior_ = 1.3 to promote oscillations and *a* = 1 to promote steep sigmoids. These two modulations define an intermediate region where *k*_1_ = 2.3. In this region, cells are in the self-organizing excitable diffusion-driven behavior presented in Figure 3D and Movie 3, giving the name to the model: the Clock and Wavefront Self-Organizing model (CWS) (Figure 4K-M and Movie 8). In this simple formulation, the model considers no posterior graded signals with PG= 0.

The CWS model forms sequential somite patterns characterized by homogeneous oscillations at the posterior tailbud, as in the CW model, but remarkably generates phase waves that travel anteriorly in the middle part of the tail in the absence of frequency gradients (white arrows Figure 4K). In the case of the CG model, local oscillator coupling generates rotating waves from random initial conditions but gives rise to coordinated anterior posterior waves within a growing tail simulation [51]. Similarly, in the CWS model, local coupling between excitable oscillators with *k*_1_ = 2.3 could generate periodic phase waves with spirals from random initial conditions (Figure 2E), but gives rise to coordinated anterior posterior somitogenesis waves within the growing tail. This is promoted by the progressive growth of the tail and by the homogeneous oscillations at the tip of the tail, which eliminates heterogeneity in refractory time guiding the patterning in the adjacent middle section.

In summary, the patterning behavior of cells in the intermediate part of the CWS model can be described as a guided self-organizing process [30], where growth and posterior oscillation guide the self-organization of phase waves in the middle part of the tail. Simulations where the tail of the CWS model is cut into two separate parts further highlight the behavior of the system when the tailbud is isolated from the rest of the tail (Figure S11, Movie 10). In agreement with experiments [55], we observed that new phase waves continue to be formed in the posterior part, while in the anterior part the phase waves that have already formed continue propagating. This illustrates that in the CWS model, the posterior oscillations guide the alternation of phases but the propagation of waves is a completely self-organizing process.

Remarkably, if the posterior oscillatory region is expanded, the model predicts the formation of multiple phase waves in the presence of a flattened gradient, providing a possible explanation for the multiple phase waves observed upon expansion and flattening of posterior gradients in mouse mutants (Figure 4L, Movie 8) [3].

In contrast to the CG model, the CWS requires local coupling between cells to generate phase waves. Indeed, without diffusion, the model will simply behave as the classical CW model presented in Figure 4C. However, as discussed in the next section and supplementary Section S17, when we reformulate the model to consider actual physical units we find that the diffusion required by the model is relatively low, in the order of 0.05 *µ*m^2^/s.

Similarly to the case of the CG model, the local coupling between oscillations in the CWS model provides robustness to noise. In addition, thanks to the independence of the model from global gradients, the somite pattern formed by the CWS is also robust to noise applied to parameter values. Indeed, the model can be implemented with any values of 0 *< k*_1_ *<* 2.3 for the posterior oscillatory region and any values of *k*_1_ *>* 4 in the anterior region. Furthermore, the excitable behavior in the intermediate region is possible for a broad set of *k*_1_ intermediate values close to the bifurcation depending on the diffusion constant *D* (Figure S5). Finally, when the same amount of noise is added to concentration and parameters in our implementations of the PORD, CG and CWS models, the somite patterns formed by the CWS model show a higher degree of relative robustness, see Figure 4M and Figure S10.

In the next section, we discuss a possible molecular implementation for the CWS model for mouse somitogenesis and discuss how the diffusion-driven patterning regime in the middle part of the tail can provide a mechanistic explanation for the change of phase between Notch and Wnt signaling observed in mouse and the excitatory behavior of mouse PSM cells in vitro.

### Possible molecular implementations of the CWS model for mouse somitogenesis

In the previous sections, we demonstrated that the Sevilletor equations provide a minimal phenomenological framework to describe various somitogenesis models as dynamical systems. This framework led us to formulate the CWS model, an extended CW model where phase waves are formed due to an excitable behavior promoted by local cell-to-cell communication independent of global graded signals, as shown in Figure 4K-M. Graded modulation, however, can be introduced into the CWS model to further recapitulate the thinning of phase waves observed along the anterior-posterior axis in vivo (Figure S13, Movie 9).

The general aim of this coarse modeling approach was to explore how the patterning behaviors of different models can arise from the global and local synchronization of oscillations of the segmentation clock, without capturing the underlying molecular details. This is a phenomenological approach as the one used in previous studies based on coupled phase oscillators [31, 21]. The Sevilletor equations, however, can also provide insights into the minimal regulatory terms between two reactants that give rise to specific patterning behaviors. In this section, we discuss two alternative molecular interpretations for the CWS model and relate each interpretation to experimental observations.

We focus our analysis on the negative feedback between *u* and *v*, which is the core regulatory topology of the Sevilletor equations that implements a delayed negative feedback without explicitly representing delays in the equations [7].

The first molecular interpretation that we consider is that the negative feedback between *u* and *v* corresponds to a transcriptional inhibition of Notch mediated by its effectors, such as members of the Hes/Her protein family. This is a key regulatory motif of the segmentation clock in vertebrates [48, 4]. According to this hypothesis, the reactant *u* can be interpreted as Notch and the reactant *v* as a cell-autonomous Notch signaling effector such as Hes7 (Figure 4N). In this scenario, diffusion represents juxtacrine Notch signaling that mediates the coupling between cells, as reported by previous studies [12, 19]. Our analysis shows that to generate a somite every 2 to 3 hours in agreement with mouse somitogenesis [46, 29], the diffusion constant of Notch has to be in the order of 0.05 *µ*m^2^/s, which is a low diffusivity consistent with juxtacrine signaling.

The second alternative molecular interpretation that we consider is that the negative feedback corresponds to coupling between Notch and Wnt signaling, which promotes oscillations in parallel to the cell-autonomous negative feedback of Notch (Figure 4O, see also supplementary Section S5). This is motivated by the observation that during mouse somitogenesis, Notch and Wnt signaling are coordinated, showing phase-shifted oscillations on the posterior side of the tail and in-phase oscillations in the middle part [2, 46]. Furthermore, perturbation experiments show that the oscillations of Wnt and Notch are entrained, supporting a regulatory feedback between the two signaling pathways [46]. According to this hypothesis, *u* can be interpreted as Wnt and *v* as Notch (Figure 4O). A simulation of an extended model presented in supplementary Section S5 shows that the cell-autonomous delayed negative feedback of Notch can coexist with the negative feedback between Wnt and Notch driving sustained oscillations (Figure 4O). For simplicity, in the rest of the study, we consider a reduced model with only the negative feedback between Wnt and Notch, which is sufficient to drive oscillation and to couple the two signaling pathways, see Figure 4P.

Remarkably, in agreement with previous quantifications [2, 46], this molecular interpretation of the CWS model predicts that Wnt and Notch oscillate out-of-phase at the posterior tip of the tail but oscillate in-phase in the intermediate region (Figure 4P). This relative phase change is promoted by the new limit cycle in the intermediate part of the tail, which arises from the excitation of cells at the two opposite bistable states, where Wnt and Notch are in phase at either high or low values. This is possible only when Wnt has a positive influence on Notch, and Notch has a negative influence on Wnt, as inverting the signs of these two cross-regulatory terms generates phase waves with out-of-phase oscillation of Wnt and Notch (supplementary Section S15). The relative change in phase between Wnt and Notch in the intermediate region can also be observed in the version of the CWS with graded modulation of *k*_1_, where, in agreement with the quantifications presented in [46], the phase waves of Notch propagate faster along the anterior-posterior axis ending up in phase with the waves of Wnt (Figure S13, Movie 9).

### The excitability of the CWS model recapitulates the behavior of the mouse PSM in-vitro

In the molecular implementation of the CWS model based on Wnt and Notch, phase wave formation is mediated by Wnt signaling. Upon rescaling the model to realistic physical units, we find that Wnt diffusion must be in the order of 0.05 *µm*^2^*/s* to generate somites every 2 or 3 hours (see Supplementary Section S17). This aligns with the low-diffusing paracrine signal of Wnt [16, 22]. Importantly, the model can incorporate Notch diffusion without altering somitogenesis patterning dynamics (see Figure S12).

As mentioned earlier, in Sevilletor equations, modulating the strength of *k*_1_ can regionalize the tail into parts with distinct dynamical regimes. In the Wnt and Notch implementation of the CWS model, this interaction corresponds to the strength of Wnt self-regulation. A plausible biological interpretation for this modulation is that other signaling pathways, such as Fgf, interact with Wnt, modulating its self-regulation along the anterior-posterior axis due to their known cross-talk during embryonic development [47].

Remarkably, two-cell simulations of cells in the novel intermediate dynamical regime of the CWS model can recapitulate the excitability of mouse PSM observed in-vitro. A previous study demonstrated that mouse PSM cells stop oscillating at low density in-vitro cultures but can be excited to oscillate when the density is increased [18]. The diffusion-driven excitable behavior of the CWS model provides a mechanistic explanation for this behavior. Indeed we found that increasing the distance between cells beyond 60 *µ*m leads to spontaneous oscillation arrest without changing any parameters (Figure 5A). The CWS model displays identical phase portraits in both high and low cell density situations, featuring two bi-stable states near unstable points (see red and yellow points in Figure 5B). However, as the distance between cells increases, weaker diffusion effects fail to push Wnt and Notch out of bi-stability toward the trajectory of the nearest unstable points explaining the arrest of oscillation (see green arrows in Figure 5B).

**Figure 5:**
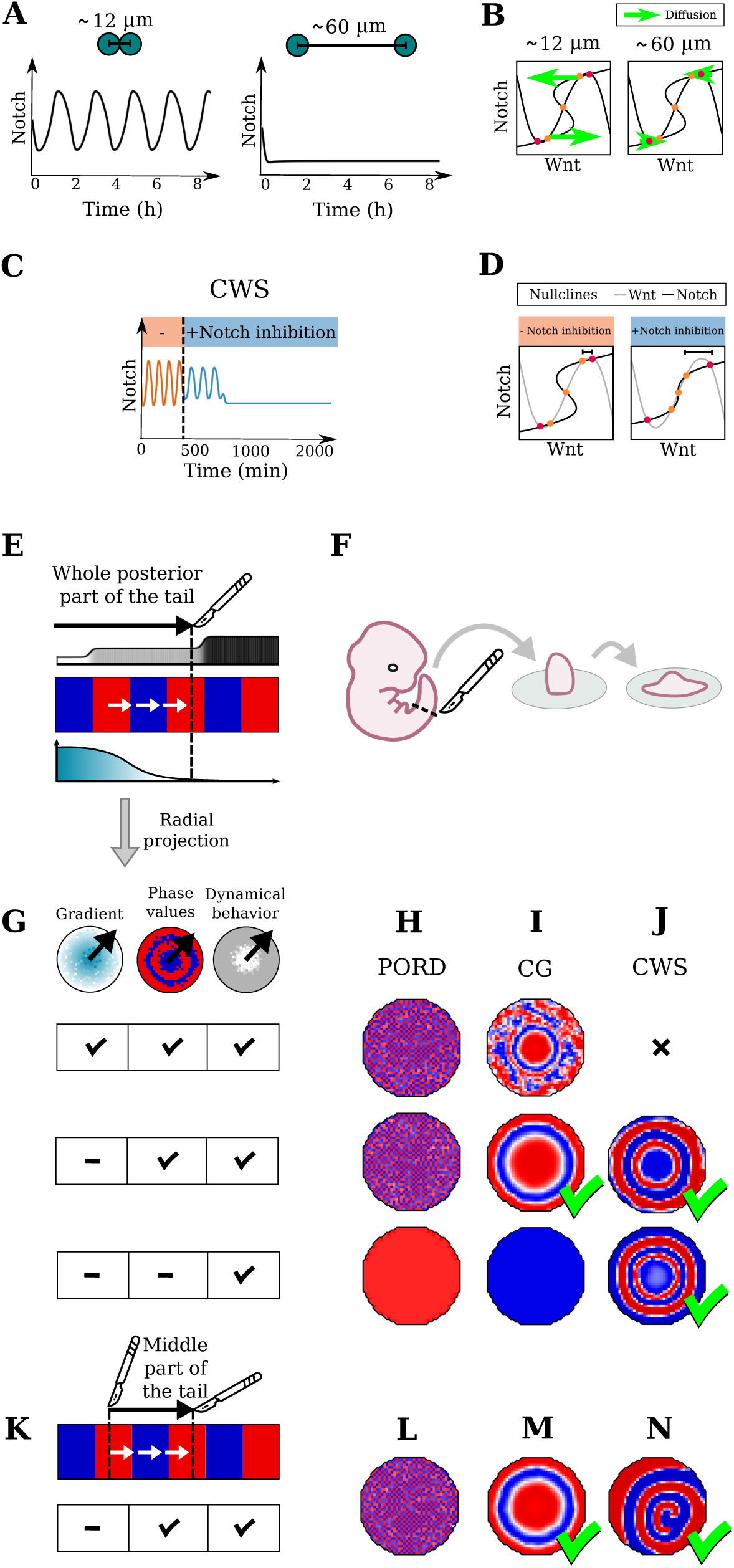
The CWS model recapitulates excitable behavior and phase waves exhibited by mouse PSM cells in-vitro. **A)** Simulation of two cells in the CWS excitable regime showing that cells are excited to oscillate when they are in close proximity (12 *µm*) and stop oscillating when they are separated (60 *µm*) similar to low-density experiments in [18] (units estimated in supplementary Section S17). **B)** Phase space shows that when cells are in close proximity they excite each other to oscillate due to the contribution of diffusion (green arrows), which decreases when cells are separated, leaving each cell at the stable states (red dots). **C)** A 2-cell simulation with the CWS excitable behavior shows that oscillations are dampened and stopped by inhibiting Notch, setting *k*_2_ = 0, similar to results shown in [18]. **D)** Without Notch inhibition in the CWS model (*k*_2_ = 1, left), the distance between the outer steady states is small. Adding Notch inhibition (*k*_2_ = 0, right) changes the shape of the nullcline of Notch (black line), increasing the distance between the outer steady states and lowering the excitability of cells. **E)** Virtual explant where the posterior portion of a simulated tail (black arrow) is projected radially and cells are shuffled to mimic cell mixing. **F)** Illustration of explant experiment. **G)** In virtual explants cells retain the specific dynamical regime (*k*_1_) of each model (gray circle), and they can retain or lose the frequency gradient (*k*_3_ and *k*_4_) (blue circle) as well as the phase values (*u* and *v* concentration) (blue and red circle). **H-J)** *v* signaling in virtual explants of the entire posterior tail with decreasing amounts of guiding cues (corresponding plot of *u* in Figure S20). Check-marks indicate whether the simulations match experimental data. **H)** In PORD explants with the gradient and/or phase values, a lateral inhibition chessboard pattern is formed due to cell mixing. Without any information, the explants form homogeneous oscillations (third row). None of the patterns match experimental findings. **I)** In CG explants, when cells inherit gradient information and phase values, salt-and-pepper oscillatory patterns are generated due to local cell rearrangement (first row). Without gradient information, cells oscillate with the same frequency according to the phase inherited from the tail generating target patterns (second row). Without any information CG explants generate homogeneous oscillations (third row). **J)** The CWS model does not have frequency gradients (see Figure 4K) so cells can retain only phase values. In this case (second row), explants generate robust target patterns with waves that propagate due to the self-organizing excitable behavior (Movie 12). The model can generate phase waves even when cells inherit only the dynamical regime, thanks to an oscillating central population of cells that guide the self-organizing process (third row). **K)** Illustration of the middle section of the virtual tail used to create explants in (L-N). **L)** Middle explants of the PORD model generate lateral inhibition chessboard patterns. **M)** Middle explants of the CG model generate target patterns because cells continue to oscillate with the initial phase values. **N)** Middle explants of the CWS model can generate target-like wave patterns promoted by an excitable bistable oscillatory regime triggered by the heterogeneity of initial phase values (Movie 11).

Our analysis highlights that excitability in the CWS model is determined by the relative magnitude between diffusion and the distance between stable and unstable steady states in the phase space. Theoretical analysis presented in Figure 3B-F demonstrates that diffusion-driven oscillations stop when diffusion remains constant but the distance between stable and unstable states increases. Perturbation experiments with mouse PSM cells in-vitro showed that inhibiting Notch signaling arrests oscillations even in high density cultures [18]. This suggests that inhibiting Notch signaling may increase the excitability threshold of cells arresting oscillations [18]. In the CWS model implemented by Wnt and Notch, the self-enhancement of Notch can be interpreted as positive feedback in response to autocrine Notch signaling [5]. Inhibition of Notch signaling therefore could correspond to the inhibition of Notch self-enhancement. Notably, Notch self-enhancement inhibition in the model leads to oscillation arrest even at high density (Figure 5C) by increasing the distance between stable and unstable steady states, elevating the excitability threshold of cells (Figure 5D).

Next, we explore how local cell communications contribute to generating coherent somitogenesis wave patterns in virtual explants of the PORD, CG, and CWS models.

### Explants of the PORD, CG and CWS model

We conducted virtual explant simulations of the somitogenesis models with decreasing levels of global information (Figure 5G). Methodological details regarding virtual explant simulations, mirroring actual explant experiments, are presented in the methods section and supplementary Section S11. Our simulations demonstrate that in PORD model explants, inheriting phase values from the tail results in the formation of chessboard patterns due to the cells’ intrinsic tendency to generate lateral inhibition patterns (Figure 5H,L and Figure S14B). This lateral inhibition behavior persists even when cells retain both gradients and phase values (first row Figure 5H), with the exception being explants that do not inherit any information from the tail, where cells undergo homogeneous oscillations (bottom row Figure 5H).

In CG model explants (Figure 5I,M and Figure S14C), we observed that cells can form target patterns when inheriting phase values from the tail but losing positional information provided by the frequency gradient (second row Figure 5I). However, inheriting positional information interferes with target pattern formation due to differences in frequencies and cell rearrangements (first row Figure 5I). Explants without information inheritance from the tail exhibit homogeneous oscillations similar to the PORD case (bottom row Figure 5I). Absence of local diffusion between cells in the CG model results in qualitatively similar patterns that are more disorganized, as cells lack local synchronization, and wave patterns are determined solely by initial phase values from the tail.

In the simplified CWS model formulation, cells lack frequency gradients (Figure 4K), and can only inherit phase values in virtual explants. In this scenario, cells generate robust target patterns characterized by circular waves propagating from the center to the periphery of the explant (second row of Figure 5J and Movie 12). This self-organizing behavior, driven by excitability, does not rely on global gradients or the initial distribution of phase values. Explants where cells do not inherit phase values (bottom row of Figure 5J) can generate circular wave patterns from homogeneous initial conditions. In this case, the formation of target patterns is guided by a central cell population acting as a “pacemaker”, reducing heterogeneties in refractory time to stimulate circular wave formation. Ablating this central population, as in experiments [18], results in either dissolving waves into homogeneous patterns or evolving into disorganized periodic phase waves depending on the initial distribution of phase values (Figure S16). A similar behavior is observed in explants derived solely from the middle part of the tail, where all cells are in the excitable regime (Figure 5N, Figure S17). These explants can produce self-organizing phase waves independent of a central cell population (Movie 11).

To further test the self-organizing capacity of the CG and CWS models we randomize cell positions to disrupt any pre-pattern inherited from the tail. Similar experiments [49, 18] have been performed by centrifuging cells from tails of different mouse embryos. In the CG model without diffusion, cells oscillate individually in a salt-and-pepper formation, while including diffusion allows local synchronization, creating patterns of phase vortices (Figure S15C,D, Movie 13). Similarly, mixed CWS model explants with diffusion exhibit periodic wave patterns generated by the excitable behavior of cells, forming rotating wave patterns (Figure S15E, Movie 13) as in the simulation with random initial conditions shown in Figure 2E with *k*_1_ = 2.3.

In summary, CWS and CG model explants intrinsically self-organize to generate periodic target wave patterns guided by inherited phase values from the tail. CWS explants additionally generate circular wave patterns from homogeneous initial phase values. In mixed explants, both models produce rotating waves, aligning with experiments [18]. It remains unclear whether rotating patterns arise from local synchronization of phases as in the CG model or as rotating spiral patterns like in the CWS model. To address this question, future explant experiments should maximize initial cell heterogeneity by mixing the intermediate part of multiple tails and remove potential guiding cues utilizing non-adherent cultures to eliminate external influences covering the effect of local synchronizations.

## Discussion

During embryonic development, cells need to coordinate their behaviors to form coherent spatial patterns that drive tissue specification. One way to achieve this coordination is by responding to global signals, such as morphogen gradients that provide positional information to the cells [53]. Alternatively, self-organizing spatial patterns can be formed by coupling cell-autonomous behaviors at the tissue level through local cell communication [50]. In addition, increasing evidence is showing that these two patterning strategies are not exclusive and that embryonic development is often controlled by self-organizing processes guided by external global signals [30].

The sequential waves of gene expression observed during vertebrate somitogenesis are a striking example of this coordination that arise from the spatial synchronizations of genetic oscillations. In this study, we introduced a system of equations named Sevilletor to investigate how oscillations can be synchronized by global signals or local cell to cell communication. We demonstrate that this system offers a minimal phenomenological framework to compare different somitogenesis patterning models, such as the Progressive Oscillatory Reaction-Diffusion (PORD) [9] and the Clock and Gradient (CG) model [31, 17, 21]. The goal of our analysis was not to reproduce the quantitative details of these models, but rather to compare the qualitative behaviors that underlie the emergence of oscillations, the formation of phase waves and arrest of oscillations in different somitogenesis models within a dynamical system.

Motivated by the observation that in the PORD and CG models the patterning of phase waves is tightly coupled with the global frequency profile, we explored whether alternative dynamical regimes of the Sevilletor model could provide additional patterning robustness by promoting phase wave formation independently of global signals.

A novel patterning behavior observed in the Sevilletor equations was the formation of periodic wave patterns. We found that this patterning regime arises near the bifurcation point between oscillation and bi-stability, where cells can be excited to oscillate by diffusion. Since in our CW implementation, posterior cells were in an oscillatory regime and anterior cells in a bi-stable regime, we envisioned an extended CW model where intermediate cells are in the novel self-organizing excitable regime. Remarkably, cells in the intermediate region were able to sustain and propagate phase waves independently of global frequency gradients, solely relying on local cell interactions. For this reason, upon the ectopic expansion of the posterior region the model predicted the formation of multiple phase waves, in agreement with experiments in mouse [3]. Our simulations revealed that this extended model has a general good robustness, even when noise is applied to the parameters that control the frequency of oscillations. We named this model the Clock and Wavefront Self-Organizing model, to highlight the hypothesis that intermediate cells are in a self-organizing regime.

While the primary focus of this study is not to uncover the molecular mechanisms underlying somitogenesis, the Sevilletor equations can offer mechanistic insights into the onset and cessation of oscillations. This led us to explore possible alternative molecular interpretations for the essential regulatory terms in the CWS model, which are discussed in more detail in the results section. Specifically, we focused our analysis on the negative feedback between two genes, which is the core regulatory term of the CWS model that drives sustained oscillations. One of the possible molecular interpretations for this negative feedback is that it corresponds to a coupling between Notch and another oscillatory pathway, such as Wnt, acting in parallel to the cell-autonomous negative feedback of Notch. This hypothesis is supported by experiments presented in [46], which agree with the prediction of the model showing that Notch and Wnt oscillate out of phase in the posterior part of the tail, and in phase in the middle part of the tail. This change of phase is promoted by the self-organizing excitable behavior of the CWS model and correlates with faster propagation of Notch signaling waves as observed experimentally [46], see Figure S13. Further supporting this molecular interpretation is the observation that Notch signaling inhibition in mouse PSM cultures reduces the excitability of cells [18], which in the CWS model is explained by reducing the self-enhancement of Notch, which can be interpreted as positive feedback in response to autocrine Notch signaling [5]. This corresponds to an increase in the distance between stable and unstable fixed points in the bistable excitable regime upon Notch signaling inhibition.

Additionally, the CWS model provides a mechanistic explanation for the observation that PSM cells stop oscillating when they are isolated in-vitro but can be excited to oscillate in higher density cultures [18]. Previously, this behavior was recapitulated with single cell simulations using the FitzHugh-Nagumo equations, where the behavior of isolated cells and high-density cultures was modeled by changing a parameter that promoted a bifurcation from bistability to oscillation [18]. In the CWS model, the transition from a quiescent state to an oscillatory state occurs with the same parameters starting from the same bi-stable regime, and is genuinely promoted by the stronger contribution of cell communication at high density cultures.

It must be noted that these predictions cannot be generalized to explain the behavior of isolated PSM Zebrafish cells in vitro, which in a recent study have been shown to oscillate autonomously for a number of cycles that depend on their anterior-posterior position in the tail [42]. Explaining the difference between the default behavior of isolated cells in mouse and Zebrafish goes beyond the scope of this study, which is mainly focused on mouse somitogenesis. Nevertheless, the broad spectrum of dynamic behaviors exhibited by the Sevilletor equations may offer a minimal framework to compare the distinct patterning regimes of different vertebrate systems. One possibility is that the difference between Zebrafish and mouse could be driven by a de-novo coupling between the autonomous oscillations of Notch and Wnt, which is not observed in Zebrafish and might modulate the default behavior of isolated PSM cells. Future theoretical models that include both feedbacks could help to explore this idea, and investigate to which extent phase waves can be formed by a cell-autonomous slowing in frequency and arrest as observed in Zebrafish [42].

Finally, to further assess the self-organizing capabilities of the CWS model and previous somitogenesis patterning hypotheses, we conducted virtual explant simulations. These simulations aimed to examine the behavior of PSM cells outside the embryonic context in each model. These experiments were motivated by previous observations demonstrating that mouse PSM explants can generate circular wave patterns despite substantial cell mixing and lack of the typical global signals in the tail [25, 18]. We found that both the CG and the CWS model are capable of generating phase waves in the absence of gradients when they inherit the phase values possessed in the tail and that the CWS model can form coherent phase waves also with initially homogeneous phase values. In addition, both models generate rotatory wave patterns in mixed explants, see Figure S15 and Movie 13. Intriguingly, similar rotating waves can be observed in explants obtained by mixing cells from different tails [18]. This behavior has also been observed in a recent model of Zebrafish somitogenesis starting from cells with randomized phases [51]. In these models the rotating waves become evident starting from more heterogeneous initial conditions. This suggests that the self-organizing behavior of the PSM can be hidden by specific initial conditions and global guiding signals, addressing the concern that the PSM lacks the characteristic features of excitable media [36]. Therefore, future experiments that aim to further reveal the self-organizing dynamics of the PSM and distinguish the prediction of different models should maximize the mixing of cells and should be based on non-adherent cultures to reduce the influence of external guides to a minimum.

In summary, the CWS model extends the CW model by considering the possibility that phase waves are formed by a self-organizing diffusion-driven excitable behavior, which is independent of global signals. This hypothesis can recapitulate four phenomena that have been observed during mouse somitogenesis. First, it accounts for the formation of multiple phase waves upon ectopic expansion and saturation of posterior gradients [3]. Secondly, it can explain the change in relative phase between Wnt and Notch observed in the posterior and middle part of the mouse tail [3]. Third, it provides a mechanistic theoretical explanation for the excitability triggered by local cell communication observed in mouse PSM cells [18]. Lastly, it elucidates how PSM cells could form circular phase wave patterns in the absence of global pre-patterns in tail explants [18].

In a broader context, the Sevilletor model offers a minimal theoretical framework for studying multi-cellular pattern formation through synchronized oscillations. In this study, we applied this framework to investigate mouse somitogenesis recapitulating the behavior of the PSM both in-vivo and in-vitro as a guided self-organizing process. In the future, we anticipate that a similar approach could be employed to study other self-organizing patterning processes during embryonic development.

## Acknowledgments

This work is supported by grant PGC2018-095170-A-100 from the Spanish Ministry of Science and Innovation and J.K. is supported by a doctoral fellowship PRE2019-087911, from the Spanish Ministry of Science and Innovation, Spain. We thank Dr. Miki Ebisuya for providing feedback.

## Contribution

Conceptualization: L.M. and J.K.; methodology: J.K. and L.M.; software: J.K.; validation: J.K. and L.M.; formal analysis: J.K. and L.M.; investigation: J.K.; data curation: J.K.; writing – original draft: J.K. and L.M, writing – review and editing: L.M. and J.K.; visualization: J.K.; supervision: L.M.; project administration: L.M.; funding acquisition: L.M.;

## Data availability

The data and codes, and any additional information required to reanalyze the data reported in this paper is available from the lead contact upon request.

## Supplementary

The supplementary materials are provided in a separate pdf file.

## Methods

### Theoretical analysis of the Sevilletor system

We devise the Sevilletor equations as a minimal reaction-diffusion system to study how genetic oscillations can synchronize and self-organize in space through local cell-to-cell communication (Figure 2A-B). The following three subsections are dedicated to a theoretical study of the properties of the network, and assumes that the reader is familiar with some aspects of complex systems theory, such as the notion of steady states, stability analysis, bifurcation diagrams and phase portraits [32, 45]

The system consists of two partial differential equations representing interactions between two reactants named *u* and *v* (equations (1) and (2), repeated below for convenience).

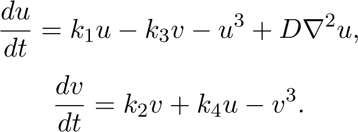

The dynamics of the system are centered around the fixed point (*u^∗^, v^∗^*) = (0, 0), which represents intermediate concentrations. However, the equations can be easily adjusted to form only positive values without affecting the behavior of the system (see supplementary material Section S1.1 and Figure S1).

The core of the system is a negative feedback between the two reactants *u* and *v*, controlled by the rates *k*_3_ and *k*_4_, which gives rise to a limit cycle centered on (*u^∗^, v^∗^*) that promotes oscillations. This is the simplest regulatory logic that can generate oscillations without explicitly adding delays to equations [7]. The sustained oscillations generated by this model are equivalent to the one generated by a single reactant model with a delayed negative feedback through a sufficiently large monotonically decreasing function [26].

In addition, the model considers cubic saturation terms that limit the deviation of concentration from the fixed point (*u^∗^, v^∗^*). These negative saturation terms should not be interpreted as degradation terms, but rather as an effective symmetric saturation for concentrations that are far from the fixed point but without significant effects for concentrations closer to the fixed point.

Finally, the model includes two positive self-regulatory feedbacks for each reactant, controlled by *k*_1_ and *k*_2_, which together with saturations determine the number of fixed points of the system. Overall, the Sevilletor equations can be viewed as an extension of the first-order formulation of the Van Der Pol oscillator [14] (supplementary Section S2) with an additional linear self-enhancing feedback and cubic saturation term in equation (2).

To characterize the behavior of the system, we focus our analysis on the effect of the positive feedbacks *k*_1_ and *k*_2_, as these are the two key parameters that drive bifurcations. Unless otherwise mentioned, we consider the non-dimensionalized version of the model for *k*_2_ by rewriting the system as *k*_2_ *→* 1, *k*_1_*/k*_2_ *→ k*_1_, *k*_3_*/k*_2_ *→ k*_3_ and *k*_4_*/k*_2_ *→ k*_4_. Importantly, for each set of reaction parameters, we also investigate the behavior of the system in the presence of diffusion by allowing the reactant *u* to diffuse with diffusion constant *D* = 0.3. For simplicity, we consider the reactant *v* to be immobile, as spatial coupling with the diffusion of one reactant (*u*) is enough to synchronize oscillations in space and to promote self-organizing patterning behaviors [41, 35, 13, 9]. In our full somitogenesis model equations (3) and (4), however, we also explore the case where *v* diffuses and find equivalent theoretical predictions.

#### Self-enhancement strength and initial conditions determine Sevilletor patterning dynamics

The bifurcation diagram in Figure 2C shows how *k*_1_ affects the number of steady states and their stability. For all values of *k*_1_, there is an unstable steady state at (*u^∗^, v^∗^*) = (0, 0) and for *k*_1_ *≥* 2.3 the system undergoes a bifurcation that adds four additional steady states (Figure 2F). The two steady states furthest away from the center are stable, while the other three steady states are unstable, see the details in the supplementary Section S1. This bifurcation is reflected by two-dimensional numerical simulations started with homogeneous initial conditions (Figure 2D-E left), showing homogeneous synchronized oscillations for 0 *≤ k*_1_ *<* 2.3 and homogeneous static patterns for *k*_1_ *≥* 2.3 associated with bistability.

Starting from random initial conditions (Figure 2D right), however, numerical simulations show a variety of complex oscillatory and static patterns (Figure 2E right) depending on the parameter *k*_1_ (Movie 1). These include lateral inhibition patterns for *k*_1_ = 0, which are characterized by cells with alternating opposite concentrations, rotating waves when *k*_1_ = 1, periodic wave patterns for *k*_1_ = 2.3, propagating bi-stable fronts that generate homogeneous static patterns when *k*_1_ = 3 and bi-stable frozen states for *k*_1_ = 4 (Movie 1). These complex patterns arise as a consequence of the combination of reaction and diffusion in the system. Since classic phase portraits only take into account the contribution of reactions, we extend our complex system analysis to study how phase portraits change with diffusion between two cells.

#### The effect of cell-to-cell communication on patterning behaviors

We use a simplified version of the Sevilletor with only two cells to study how diffusion affects the patterning behavior. The cells start with a heterogeneous initial state with (*u*_1_*, v*_1_) = (0.1, 0) and (*u*_2_*, v*_2_) = (*−*0.1, 0). We run two simulations for each representative value of *k*_1_: one without diffusion (*D* = 0), and one with diffusion (*D* = 0.3) (Figure 3B-F). Without diffusion, each cell acts as an individual unit and its behavior depends solely on changes driven by reaction (Figure 3B-F left columns). This case corresponds to the behavior seen in square two-dimensional simulations with a homogeneous initial state (Figure 2E left column), since when every cell has the exact same amounts of *u* and *v* there is no active contribution from diffusion, i.e. *∇*^2^*u* = 0. By including cell communication in the form of diffusion of *u* (Figure 3B-F right columns), the combined effect of reaction and diffusion coordinates the behavior of the two cells, giving rise to large scale patterns.

With *k*_1_ = 0 and *k*_1_ = 1, the cells oscillate individually in the limit cycle without diffusion. For *k*_1_ = 0, with diffusion these oscillatory trajectories are counterbalanced and stabilized by the effect of diffusion (Figure 3B, Movie 2) [44]. Our 2D simulations show that this behavior generates a lateral inhibition (chessboard) pattern from noise (Figure 2E and first column in Movie 1). With *k*_1_ = 1, with diffusion the cells continue to oscillate by following a smaller limit cycle that in the long run synchronizes the two cells together (Figure 3C). In 2D simulations this behavior makes rotating waves from noise (Figure 2E, second column in Movie 1).

At the bifurcation point *k*_1_ = 2.3, in the absence of diffusion, the two cells do not oscillate and are trapped at the nearest stable state on the upper or lower half plane (Figure 3D). When diffusion is added, however, the equilibrating effect of diffusion pushes cells out of stability towards the trajectory of the closest stable state (green arrows in Figure 3D). Following this trajectory, each cell goes to the opposite half-plane towards the stable steady state, and it is again destabilized by diffusion. The repetition of this process generates a novel limit cycle that keeps cells oscillating (Movie 3). In 2D simulations with random initial conditions, this dynamics gives rise to a new type of periodic wave pattern with spiral formation that to the best of our knowledge has not been described previously (Figure 2E, third column in Movie 1) (see detailed analysis in Supplementary Section S2). The limit cycle that underlies periodic wave patterns is possible for a variety of values of *k*_1_ and diffusion constant *D* (Figure S5).

This dynamical behavior changes for larger values of *k*_1_. For example, for a self-enhancement strength with *k*_1_ = 3, a few oscillations are stimulated, but the two cells eventually freeze together at the same stable state (Figure 3E). This makes a propagating front that covers the whole domain in square 2D simulations (Figure 2E, fourth column of Movie 1). For even stronger self-enhancement with *k*_1_ = 4, the distance between stable states and saddle points is too large for diffusion to impact the trajectory of the cells (Figure 3F), making a static bi-stable frozen pattern that amplifies the pre-pattern present in the initial conditions (Figure 2E, fifth column of Movie 1).

A characteristic of the Sevilletor system over previous reaction-diffusion models is that its patterning dynamics can be changed by varying just the parameter *k*_1_. This property can also be exploited to easily switch between different patterning behaviors over time (Movie 4).

#### The relative self-enhancement strength and diffusion determines the dynamical behavior the system

To further characterize the effect of both self-enhancements, we performed a 2D simulation of the full system with equations (1) and (2) by increasing the values of *k*_1_ and *k*_2_ along the y- and x-axis to recapitulate all the different dynamical behaviors of the system within the same simulation (Figure 3H, Movie 5). The corresponding 2D bifurcation diagram (Figure 3I) shows the bifurcation between 1, 5 and 9 steady states. This bifurcation diagram was further divided into regions of different patterning behaviors by calculating the number of oscillations of a series of two-cell simulations with diffusion (as in Figure 3B-F right columns) for each combination of parameters (*k*_1_*, k*_2_) in a 100*x*100 grid for a total of 10000 two-cell simulations (Figure 3I). This allowed us to derive an extended bifurcation diagram where different colored regions correspond to the different dynamical behaviors that the system can generate with diffusion (Figure 3J). This extended bifurcation diagram shows that starting from random initial conditions the relative self-enhancement strength of *k*_1_ and *k*_2_ together with diffusion determines the different patterning behaviors of the system.

### Simulation details

Simulations are run with a finite difference solver written in Julia 1.6.4. The complex systems analysis and two cell simulations have been run in Mathematica 12, the reverse-LUT method has been written in Python 3.8.10, and Fiji (ImageJ) has been used to analyze the patterns seen in the simulated and experimental explants. Python 3.8.10 is used to create all plots, excluding the phase spaces and bifurcation diagrams which have been generated using Mathematica 12.

#### Time discretization

In all simulations, we discretize time by using an Euler method to update the values of *u* and *v* for each cell in the system with position (*x, y*):

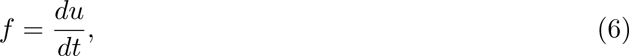

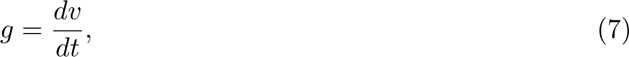

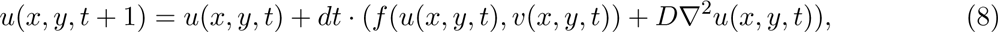

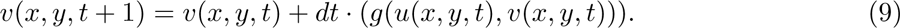

#### Space discretization

The diffusion term *∇*^2^*u*(*x, y*) in the system is calculated using a first order finite difference scheme to calculate the discrete Laplace. This is done by using a Taylor expansion of the functions *u*(*x, y, t*) and *v*(*x, y, t*) around the steady state point (0,0). *ɛ* = *dx* = *dy* in the following. For *u* the calculations are:

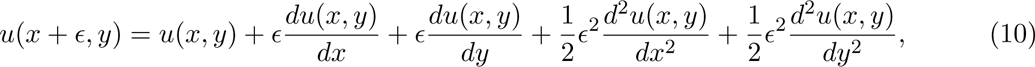

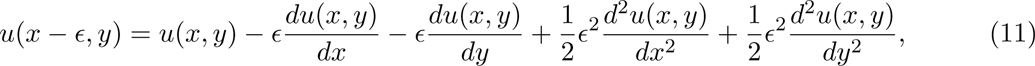

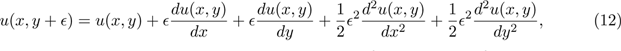

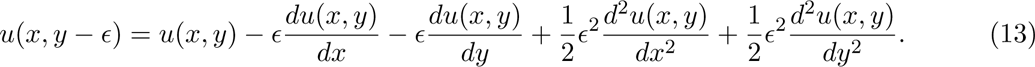

We isolate the first order differential terms in the equations and combine the equations to get:

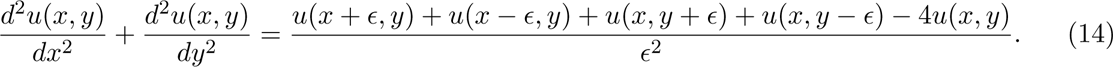

Similar calculations are also performed for *v*.

#### Boundary conditions

In all simulations, except the one in Figure S4B, we consider zero flux Neumann boundary conditions. This means that nothing diffuses between the outside and inside of the system, i.e. the Laplace is equal to zero at the boundaries. The Laplace at the boundary is derived as:

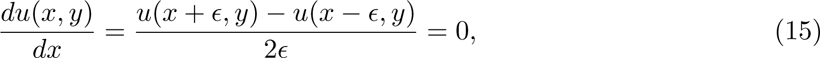

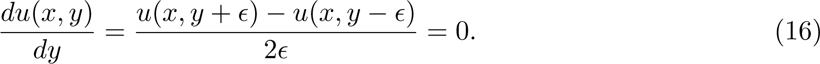

This leads to:

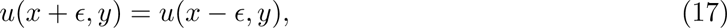

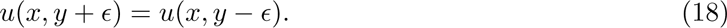

Inserting the appropriate ones for the edge/corner into the discrete Laplace equation (14) gives the correct formula, for example for the bottom left corner (*x, y*) = (1, 1) in equation (19):

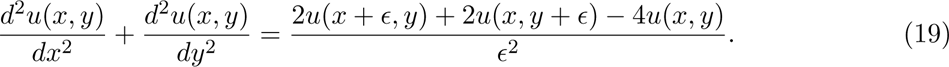

The periodic boundary conditions used for the simulation in Figure S4B are also calculated using the discrete Laplace, treating cells on the boundaries as direct neighbors with cells on the corresponding boundary. For example for the cell in the bottom left corner (*x, y*) = (1, 1) in a system where the lengths of the axes are *L*:

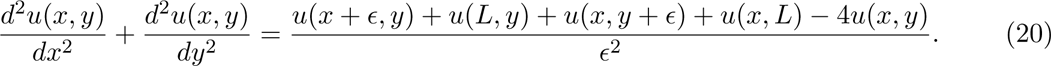

#### Parameters

- *dx* = *dy* = 1
- Diffusion constants:

**–** Square simulations in Figure 2 and 3: *D* = 0.3
**–** PORD: *D* = 1.0
**–** Clock and Wavefront model: *D* = 0
**–** Clock and Gradient model: *D* = 0.3
**–** Clock and Wavefront Self-Organizing model: *D* = 0.3 For other exceptions included in the supplementary the value of *D* is given
- For the square pattern systems in Figures 2 and 3: *dt* = 0.005. For the somitogenesis models: *dt* = 0.002
- Size of the square pattern systems in Figures 2 and 3: *L_x_* = *L_y_* = 100
- Initial size of the virtual tails

**–** PORD: *L_x_* = 50 and *L_y_* = 12
**–** Clock and Wavefront model: *L_x_* = 35 and *L_y_* = 20
**–** Clock and Gradient model: *L_x_* = 35 and *L_y_* = 20
**–** Clock and Wavefront Self-Organizing model: *L_x_*= 35 and *L_y_* = 20
- Number of cells in virtual explants

**–** PORD: *N* = 1245
**–** Clock and Gradient model: *N* = 1245
**–** Clock and Wavefront Self-Organizing model: *N* = 1245
- The frequency of adding a new row of cells during the somitogenesis simulation in a virtual tail

**–** PORD: every 1700th time step
**–** Clock and Wavefront model: every 200th time step
**–** Clock and Gradient model: every 100th time step
**–** Clock and Wavefront Self-Organizing model: every 200th time step
- Initial state of the simulation

**–** Square simulations in Figure 2 and 3: mean *u* = 0 and *v* = 0 and std=0.1 for the initial states with noise and std=0 for homogeneous initial conditions
**–** PORD: *u* = 0 and *v* = 1 in the top row and *v* = 0 everywhere else
**–** Clock and Wavefront model: *u* = *−*1 and *v* = *−*1 for all cells
**–** Clock and Gradient model: *u* = *−*1 and *v* = *−*1 for all cells
**–** Clock and Wavefront Self-Organizing model: *u* = *−*1 and *v* = *−*1 for all cells
- *k*_1_, *k*_2_ and *k*_3_:

**–** PORD: *k*_1_ = 0 for all cells and *k*_3_ and *k*_4_ is a gradient from 1.3 to 1
**–** Clock and Wavefront model: *k*_1_ = 1 on the posterior side and *k*_1_ = 4 on the anterior side. *k*_3_ = *k*_4_ = 1 for all cells
**–** Clock and Gradient model: *k*_1_ = 1 on the posterior side and *k*_1_ = 4 on the anterior side. *k*_3_ and *k*_4_ is a gradient from 3 to 1
**–** Clock and Wavefront Self-Organizing model: *k*_1_ = 1 on the posterior side, *k*_1_ = 2.3 in the middle section and *k*_1_ = 4 on the anterior side. *k*_3_ = *k*_4_ = 1 for all cells
- Multiplicative noise in the simulations in Figure 4 and Figure S10: 5% noise is added to the values of *u* and *v* and 0.5% noise is added to the parameters *k*_1_*, k*_3_, and *k*_4_ every time a new row of cells is added on the posterior side. An exception is the PORD model, where no noise is added to the value of *k*_1_ = 0.

### Quantification and statistical analysis

#### Stability and types of steady states

The steady states of the Sevilletor model and their properties are calculated with a phase plane analysis [32]. The steady states of a system are all pairs (*u^∗^, v^∗^*) for which 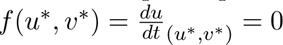 and 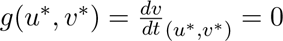. The stability of the steady states are found by using linear stability analysis from the determinant and trace of the matrix *A*:

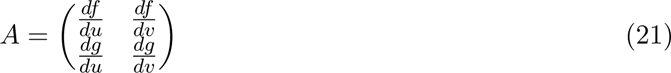

For Det(*A*_(_*_u_∗_,v_∗*_)_) *>* 0 and Trace(*A*_(_*_u_∗_,v_∗*_)_) *<* 0 the steady state is stable, otherwise it is unstable. The type of steady state is identified as discussed in Appendix A in [32] and describes the shape of the vector field around the steady state.

#### Calculation of the phase and the number of oscillations performed in Figure 3

The phase is calculated, at every time point, as the angle between the initial position and the position of the cell in phase space around the central unstable steady state in (*u, v*) = (0, 0).

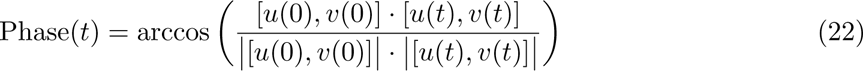

The number of oscillation is determined by counting the local minima that are close to zero along the phase profile, excluding the first minima for *t* = 0.

**Figure M1:**
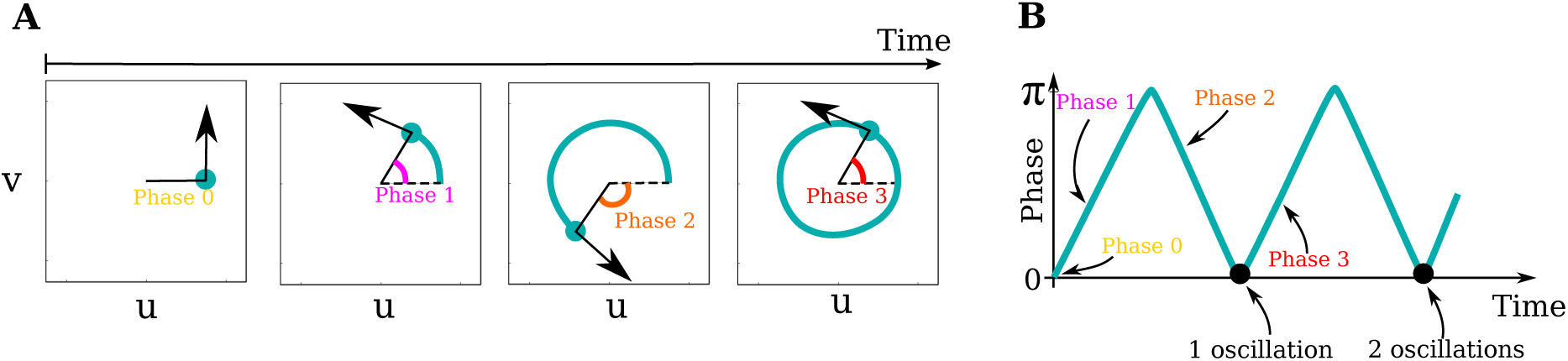
Details of the calculation of phase and number of oscillations in Figure 3. **A)** The timeline shows the path of a cell in the phase space. The phase is calculated for each time point, starting from 0, increasing to *π* for half a loop, and decreasing back to 0 for the second half of the loop. **B)** The phase is plotted as a function of time and the number of oscillations is measured as the number of local minima in phase = 0 excluding the initial for *t* = 0, marked by black points.

## References

[1] Alexander Aulehla and Olivier Pourquíe. Signaling gradients during paraxial mesoderm development. Cold Spring Harbor Perspectives in Biology, 2:a000869, 2010.

[2] Alexander Aulehla, Christian Wehrle, Beate Brand-Saberi, Rolf Kemler, Achim Gossler, Benoit Kanzler, and Bernhard G. Herrmann. Wnt3a plays a major role in the segmentation clock controlling somitogenesis. Developmental Cell, 4:395–406, 2003.

[3] Alexander Aulehla, Winfried Wiegraebe, Valerie Baubet, Matthias B. Wahl, Chuxia Deng, Makoto Taketo, Mark Lewandoski, and Olivier Pourquíe. A beta-catenin gradient links the clock and wavefront systems in mouse embryo segmentation. Nature Cell Biology, 10:186–193, 2008.

[4] Ahmet Ay, Stephan Knierer, Adriana Sperlea, Jack Holland, and Ertuğrul M. Ö zbudak. Short-lived her proteins drive robust synchronized oscillations in the zebrafish segmentation clock. Development, 140:3244–3253, 8 2013.

[5] Robert A. Bone, Charlotte S. L. Bailey, Guy Wiedermann, Zoltan Ferjentsik, Paul L. Appleton, Philip J. Murray, Miguel Maroto, and J. Kim Dale. Spatiotemporal oscillations of notch1, dll1 and nicd are coordinated across the mouse psm. Development, 141:4806–4816, 2014.

[6] Michael Breakspear, Stewart Heitmann, and Andreas Daffertshofer. Generative models of cortical oscillations: Neurobiological implications of the kuramoto model. Frontiers in Human Neuroscience, 4, 2010.

[7] Pablo Casani-Galdon and Jordi Garcia-Ojalvo. Signaling oscillations: Molecular mechanisms and functional roles. Current Opinion in Cell Biology, 78:102130, 2022.

[8] J. Cooke and E. C. Zeeman. A clock and wavefront model for control of the number of repeated structures during animal morphogenesis. Journal of Theoretical Biology, 58:455–476, 1976.

[9] James Cotterell, Alexandre Robert-Moreno, and James Sharpe. A local, self-organizing reaction-diffusion model can explain somite patterning in embryos. Cell Systems, 1:257–269, 2015.

[10] Julien Dubrulle, Michael J. McGrew, and Olivier Pourquíe. Fgf signaling controls somite boundary position and regulates segmentation clock control of spatiotemporal hox gene activation. Cell, 106:219–232, 2001.

[11] Gregg Duester. Retinoic acid regulation of the somitogenesis clock. Birth Defects Research Part C - Embryo Today: Reviews, 81:84–92, 2007.

[12] Zoltan Ferjentsik, Shinichi Hayashi, J. Kim Dale, Yasumasa Bessho, An Herreman, Bart De Strooper, Gonzalo Del Monte, Jose Luis De La Pompa, and Miguel Maroto. Notch is a critical component of the mouse somitogenesis oscillator and is essential for the formation of the somites. PLoS Genetics, 5, 9 2009.

[13] Richard J. Field and Richard M. Noyes. Oscillations in chemical systems. iv. limit cycle behavior in a model of a real chemical reaction. The Journal of Chemical Physics, 60:1877–1884, 1974.

[14] Richard FitzHugh. Impulses and physiological states in theoretical models of nerve membrane. Biophysical Journal, 1:445–466, 1961.

[15] Paul François, Vincent Hakim, and Eric D. Siggia. Deriving structure from evolution: Metazoan segmentation. Molecular Systems Biology, 3:154, 2007.

[16] Yudai Hatakeyama, Nen Saito, Yusuke Mii, Ritsuko Takada, Takuma Shinozuka, Tatsuya Takemoto, Honda Naoki, and Shinji Takada. Intercellular exchange of wnt ligands reduces cell population heterogeneity during embryogenesis. Nature Communications, 14, 12 2023.

[17] Leah Herrgen, Saúl Ares, Luis G. Morelli, Christian Schröter, Frank Jülicher, and Andrew C. Oates. Intercellular coupling regulates the period of the segmentation clock. Current Biology, 20:1244–1253, 2010.

[18] Alexis Hubaud, Ido Regev, L. Mahadevan, and Olivier Pourquíe. Excitable dynamics and yap-dependent mechanical cues drive the segmentation clock. Cell, 171:668–682.e11, 2017.

[19] Yun-Jin Jiang, Birgit L. Aerne, Lucy Smithers, Catherine Haddon, David Ish-Horowicz, and Julian Lewis. Notch signalling and the synchronization of the somite segmentation clock. Nature, 408:475–479, 11 2000.

[20] Laurent Jutras-Dubé, Ezzat El-Sherif, and Paul François. Geometric models for robust encoding of dynamical information into embryonic patterns. eLife, 9:e55778, 2020.

[21] David J. Jörg, Andrew C. Oates, and Frank Jülicher. Sequential pattern formation governed by signaling gradients. 10 2016.

[22] Anna Kicheva, Periklis Pantazis, Tobias Bollenbach, Yannis Kalaidzidis, Thomas Bitting, Thomas Jülicher, and Marcos Gonźalez-Gaitάn. Kinetics of morphogen gradient formation. Science, 315, 1 2007.

[23] Yoshiki Kuramoto. *Self-entrainment of a population of coupled non-linear oscillators*, pages 420–422. Springer-Verlag, 1975.

[24] Chandrashekar Kuyyamudi, Shakti N. Menon, and Sitabhra Sinha. Morphogen-regulated contact-mediated signaling between cells can drive the transitions underlying body segmentation in vertebrates. Physical Biology, 19:16001, 2022.

[25] Volker M. Lauschke, Charisios D. Tsiairis, Paul Fraņcois, and Alexander Aulehla. Scaling of embryonic patterning based on phase-gradient encoding. Nature, 493:101–105, 2013.

[26] Julian Lewis. Autoinhibition with transcriptional delay: A simple mechanism for the zebrafish somitogenesis oscillator. Current Biology, 13:1398–1408, 2003.

[27] Maroto Maroto, J. Kim Dale, Mary Lee Deqúeant, Anne Ćecile Petit, and Olivier Pourquíe. Synchronised cycling gene oscillations in presomitic mesoderm cells require cell-cell contact. International Journal of Developmental Biology, 49:309–315, 2005.

[28] Yoshito Masamizu, Toshiyuki Ohtsuka, Yoshiki Takashima, Hiroki Nagahara, Yoshiko Takenaka, Kenichi Yoshikawa, Hitoshi Okamura, and Ryoichiro Kageyama. Real-time imaging of the somite segmentation clock: Revelation of unstable oscillators in the individual presomitic mesoderm cells. PNAS, 103:1313–1318, 2006.

[29] Mitsuhiro Matsuda, Hanako Hayashi, Jordi Garcia-Ojalvo, Kumiko Yoshioka-Kobayashi, Ryoichiro Kageyama, Yoshihiro Yamanaka, Makoto Ikeya, Junya Toguchida, Cantas Alev, and Miki Ebisuya. Species-specific segmentation clock periods are due to differential biochemical reaction speeds. Science, 369:1450–1455, 2020.

[30] J. Serrano Morales, Jelena Raspopovic, and Luciano Marcon. From embryos to embryoids: How external signals and self-organization drive embryonic development. Stem Cell Reports, 16:1039–1050, 2021.

[31] Luis G. Morelli, Saúl Ares, Leah Herrgen, Christian Schröter, Frank Jülicher, and Andrew C. Oates. Delayed coupling theory of vertebrate segmentation. HFSP Journal, 3:55–66, 2009.

[32] J. D. Murray. Mathematical Biology I. An Introduction. Springer, 3rd edition, 2002.

[33] L. A. Naiche, Nakisha Holder, and Mark Lewandoski. Fgf4 and fgf8 comprise the wavefront activity that controls somitogenesis. PNAS, 108:4018–4023, 2011.

[34] Stuart A. Newman. Self-organization in embryonic development: myth and reality. Preprint in arxiv.org, 2022.

[35] G. Nicolis and I. Prigogine. Self-Organization in Nonequilibrium Systems. From Dissipative Structures to Order through Fluctuations. Wiley-Interscience, 1977.

[36] Andrew C. Oates. Waiting on the fringe: cell autonomy and signaling delays in segmentation clocks. Current Opinion in Genetics and Development, 63:61–70, 2020.

[37] Andrew C. Oates, Luis G. Morelli, and Saúl Ares. Patterning embryos with oscillations: Structure, function and dynamics of the vertebrate segmentation clock. Development, 139:625–639, 2012.

[38] Isabel Palmeirim, Domingos Henrique, David Ish-Horowicz, and Olivier Pourquíe. Avian hairy gene expression identifies a molecular clock linked to vertebrate segmentation and somitogenesis. Cell, 91:639–648, 1997.

[39] Jasmina Panovska-Griffiths, Karen M. Page, and James Briscoe. A gene regulatory motif that generates oscillatory or multiway switch outputs. Journal of the Royal Society Interface, 10, 2 2013.

[40] Jesús Pantoja-Herńandez, Víctor F. Breña-Medina, and Moiśes Santilĺan. Hybrid reaction-diffusion and clock-and-wavefront model for the arrest of oscillations in the somitogenesis segmentation clock. Chaos: An Interdisciplinary Journal of Nonlinear Science, 31:063107, 2021.

[41] I. Prigogine and R. Lefever. Symmetry breaking instabilities in dissipative systems. ii. The Journal of Chemical Physics, 48:1695–1700, 1968.

[42] Laurel A. Rohde, Arianne Bercowsky-Rama, Jose Negrete Jr, Guillaume Valentin, Sundar Ram Naganathan, Ravi A. Desai, Petr Strnad, Daniele Soroldoni, Frank Jülicher, and Andrew C. Oates. Cell-autonomous generation of the wave pattern within the vertebrate segmentation clock. Preprint in bioRxiv, 2021.

[43] I. A. Shepelev and T. E. Vadivasova. Variety of spatio-temporal regimes in a 2d lattice of coupled bistable fitzhugh-nagumo oscillators. formation mechanisms of spiral and double-well chimeras. Communications in Nonlinear Science and Numerical Simulation, 79:104925, 2019.

[44] Rajeev Singh and Sitabhra Sinha. Spatiotemporal order, disorder, and propagating defects in homogeneous system of relaxation oscillators. *Physical Review E - Statistical*, Nonlinear, and Soft Matter Physics, 87:012907, 2013.

[45] Kim Sneppen. Models of Life - Dynamics and Regulation in Biological Systems. Cambridge University Press, 1st edition, 2014.

[46] Katharina F. Sonnen, Volker M. Lauschke, Julia Uraji, Henning J. Falk, Yvonne Petersen, Maja C. Funk, Mathias Beaupeux, Paul François, Christoph A. Merten, and Alexander Aulehla. Modulation of phase shift between wnt and notch signaling oscillations controls mesoderm segmentation. Cell, 172:1079–1090.e12, 2018.

[47] Michael J. Stulberg, Aiping Lin, Hongyu Zhao, and Scott A. Holley. Crosstalk between fgf and wnt signaling in the zebrafish tailbud. Developmental Biology, 369:298–307, 2012.

[48] Yoshiki Takashima, Toshiyuki Ohtsuka, Aitor Gonźalez, Hitoshi Miyachi, and Ryoichiro Kageyama. Intronic delay is essential for oscillatory expression in the segmentation clock. PNAS, 108:3300–3305, 2 2011.

[49] Charisios D. Tsiairis and Alexander Aulehla. Self-organization of embryonic genetic oscillators into spatiotemporal wave patterns. Cell, 164:656–667, 2016. Somitogenesis.

[50] A. M. Turing. The chemical basis of morphogenesis. Philos. Trans. R. Soc. Lond. B, 237:37–72, 1952.

[51] Koichiro Uriu, Bo Kai Liao, Andrew C. Oates, and Luis G. Morelli. From local resynchronization to global pattern recovery in the zebrafish segmentation clock. eLife, 10, 2 2021.

[52] Alexis B. Webb, Iván M. Lengyel, David J. Jörg, Guillaume Valentin, Frank Jülicher, Luis G. Morelli, and Andrew C. Oates. Persistence, period and precision of autonomous cellular oscillators from the zebrafish segmentation clock. eLife, 5:e08438, 2016.

[53] L. Wolpert. Positional information and the spatial pattern of cellular differentiation. Journal of Theoretical Biology, 25:1–47, 1969.

[54] A. M. Zhabotinsky and A. N. Zaikin. Autowave processes in a distributed chemical system. Journal of Theoretical Biology, 40:45–61, 1973.

[55] Ece Ö zeļci, Erik Mailand, Matthias Rüegg, Andrew C. Oates, and Mahmut Selman Sakar. Deconstructing body axis morphogenesis in zebrafish embryos using robot-assisted tissue micromanipulation. Nature Communications, 13, 12 2022.

