## Supplementary materials for "A Clock and Wavefront Self-Organizing Model Recreates the Dynamics of Mouse Somitogenesis in-vivo and in-vitro"

PREPRINT RESEARCH ARTICLE

January 17, 2024

### Contents

**S1 Complex systems analysis of the Sevilletor network** **1**
S1.1 Sevilletor with only positive values of $u$ and $v$ . . . . . . . . . . . . . . . . . . . . . . 1
S1.2 Detailed analysis of the effect of cell-to-cell communication on patterning behaviors . 1
S1.3 Dispersion relation diagrams . . . . . . . . . . . . . . . . . . . . . . . . . . . . . . . . 3

**S2 Models that generate rotating wave patterns** **4**

**S3 Parameter space that gives rise to periodic wave patterns** **7**

**S4 Initial values of u and v determine different types of spatial synchronizations** **8**

**S5 Adding a self-regulatory delayed negative feedback of Notch does not change the oscillatory behavior of Wnt and Notch** **9**

**S6 The original PORD model by Cotterell et al. [3]** **10**

**S7 The CWS model is robust to noise** **13**

**S8 Dissection of the tailbud in the CWS model** **14**

**S9 Adding diffusion of Notch to the CWS model** **15**

**S10 CWS model with thinning of the waves along the anterior-posterior axis** **16**

**S11 The CWS model recapitulates the rotating wave patterns observed in mixed explant experiments** **17**

**S12 Varying the position of the cut in explants** **18**

**S13 Ablating the center of explants from the Clock and Wavefront Self-Organizing model** **20**

**S14 Possible outcomes of middle tail explants in CWS model** **21**

**S15 Reversed negative feedback between Wnt and Notch in the CWS** **22**

**S16 Wnt patterns in models of virtual tails and explants** **23**

**S17 Units in the CWS model** **25**

**S18 Summary of Figure 4** **26**

**S19 Captions of Movies** **27**

### S1 Complex systems analysis of the Sevilletor network

#### S1.1 Sevilletor with only positive values of $u$ and $v$

The patterns generated by the Sevilletor model are centered around the fixed point $(u^*, v^*) = (0, 0)$, which represents intermediate concentrations. In this context, negative values do not represent negative concentrations but rather a decrease from intermediate values. The equations can easily modified to generate only positive values by substituting $u$ and $v$ in equations (1) and (2) as $u \rightarrow (u - c)$ and $v \rightarrow (v - c)$, as shown in equations (S1) and (S2). An example is shown in Figure S1B, where a periodic wave pattern is formed centered around the value $c = 2$ with $k_1 = 2.3$ and $k_2 = 1$.

$$\frac{du}{dt} = k_1(u - c) - (v - c) - (u - c)^3 + D\nabla^2(u - c), \tag{S1}$$

$$\frac{dv}{dt} = (v - c) + (u - c) - (v - c)^3. \tag{S2}$$

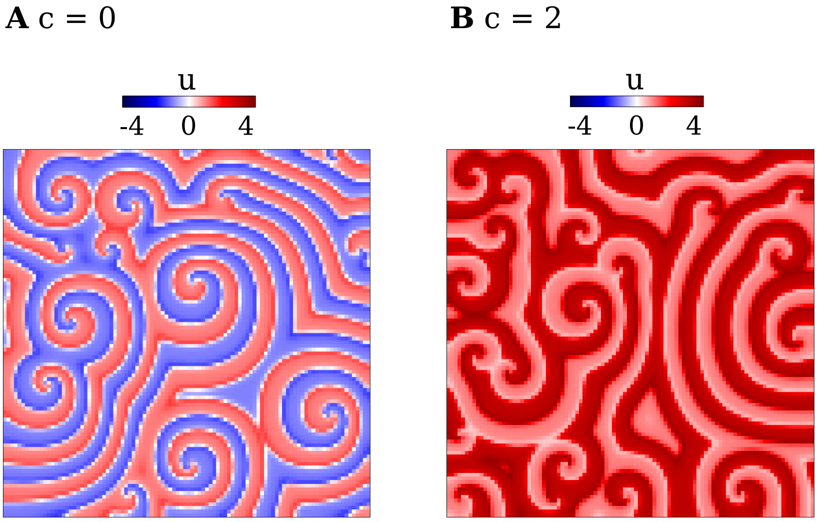


Figure S1: **Example of simulations with only positive values of u and v.**
Examples of periodic wave patterns with spirals generated by equations (S1) and (S2) where $k_1 = 2.3$ showing the values of $u$ in a simulation started from noise.
**A)** With values of $u$ and $v$ centered around 0 with $c = 0$.
**B)** With only positive values of $u$ and $v$ with $c = 2$.

#### S1.2 Detailed analysis of the effect of cell-to-cell communication on patterning behaviors

We consider a simplified version of the Sevilletor model implemented in a system with only two cells to study how diffusion affects the patterning behavior. We simulate this simplified model starting from a heterogeneous initial state with $(u_1, v_1) = (0.1, 0)$ and $(u_2, v_2) = (-0.1, 0)$ for the two cells respectively and run two simulations for each representative value of $k_1$: one without diffusion ($D = 0$), and one with diffusion ($D = 0.3$) (Figure 3B-F). Without diffusion, each cell acts as an individual unit and its behavior depends solely on changes driven by reaction (Figure 3B-F left columns). This case corresponds to the behavior seen in two-dimensional simulations with a homogeneous initial state (Figure 2E left column), since when every cell has the exact same amounts of $u$ and $v$ there is no active contribution from diffusion, i.e. $\nabla^2 u = 0$. By including cell communication in the form of diffusion of $u$ (Figure 3B-F right columns), the combined effect of reaction and diffusion coordinates the behavior of the two cells, giving rise to large scale patterns. To characterize these behaviors we also quantify phase changes over time (details of the phase calculation are provided in methods section).

With $k_1 = 0$, in the absence of diffusion, the two cells undergo continuous oscillations with independent trajectories in the limit cycle. In the presence of diffusion, however, these oscillatory trajectories are counterbalanced by the effect of diffusion, which stabilizes the two cells in a frozen out-of-phase configuration (Figure 3B, Movie 2) [16]. Our two-dimensional simulations show that this type of behavior generates a chessboard pattern where each cell has a frozen phase opposite to that of its neighbors, as in lateral inhibition patterns (right column of the first row in Figure 2E and first column in Movie 1).

When $k_1 = 1$, in the absence of diffusion, the two cells oscillate independently. When diffusion is added to the model, the cells continue to oscillate by following a smaller limit cycle that in the long run synchronizes the two cells together as in a type IIIo system [4] (Figure 3C). Using random initial concentrations that lay on one of the half planes, this behavior is associated with homogeneous oscillations (Figure S6). However, starting from random initial concentrations spread on the two half planes, the system generates rotating spirals similar to those formed by a diffusive Van der Pol oscillator (right column of the second row Figure 2E, Figure S3 and second column in Movie 1).

At the bifurcation point $k_1 = 2.3$, in the absence of diffusion, the two cells do not oscillate and are trapped at the nearest stable state on the upper or lower half plane. When diffusion is added to the model, however, the equilibrating effect of diffusion pushes cells out of stability towards the trajectory of the closest saddle point (green arrows in Figure 3D). Following this trajectory, each cell goes to the opposite half-plane towards the stable steady state, and it is again destabilized by diffusion. The repetition of this process generates a novel limit cycle that keeps cells oscillating (Movie 3). This new limit cycle generates in-phase oscillations of $u$ and $v$, because the permanence time around the stable states is greater than the time it takes to follow the trajectory to the opposite half-plane (Movie 3). In two-dimensional simulations with random initial conditions, this dynamics gives rise to a new type of spiral patterning behavior that to the best of our knowledge has not been described previously (Figure 2E third row and third column in Movie 1). The pattern looks very similar to those formed by classic models of the Belousov–Zhabotinsky reaction [22] such as the Brusselator [14] and Oregonator [5], however it emerges from a different dynamical behavior that has not been described in previous models (see detailed analysis in Supplementary Section S2).

The limit cycle that underlies stripy spiral patterns is possible for a variety of values of $k_1$ and diffusion constant $D$ (Figure S5). Given a fixed value of $D$, the dynamical behavior changes for different values of $k_1$, because the distance between the stable steady states and saddle points will change. For example, for a larger self-enhancement strength with $k_1 = 3$, the effect of diffusion is strong enough to stimulate a few oscillations, but the two cells eventually freeze together at the same stable state (Figure 3E). In two-dimensional simulations, this behavior generates a propagating front of one stable state that covers the whole domain, giving rise to a homogeneous static pattern (Figure 2E fourth row and fourth column of Movie 1). For even stronger self-enhancement with $k_1 = 4$, the distance between stable states and saddle points is too large for diffusion to impact the trajectory of the cells. Therefore, the two cells are frozen on the closest steady state (Figure 3F). In two-dimensional simulations this behavior generates a static bi-stable frozen pattern that amplifies the noise present in random initial conditions (Figure 2E fifth row and fifth column of Movie 1).

A major benefit of the Sevilletor model over previous models is that its patterning dynamics can be changed by varying just the parameter $k_1$. This property can be exploited to switch between different patterning behaviors over time, see for example the simulation in the Movie 4, where the model switches between a lateral inhibition pattern to a rotating spiral pattern, a stripy spiral pattern, a bi-stable frozen spiral pattern and finally back to lateral inhibition pattern $k_1 = 0.0 \rightarrow 1.0 \rightarrow 2.3 \rightarrow 4.0 \rightarrow 0.0$.

#### S1.3 Dispersion relation diagrams

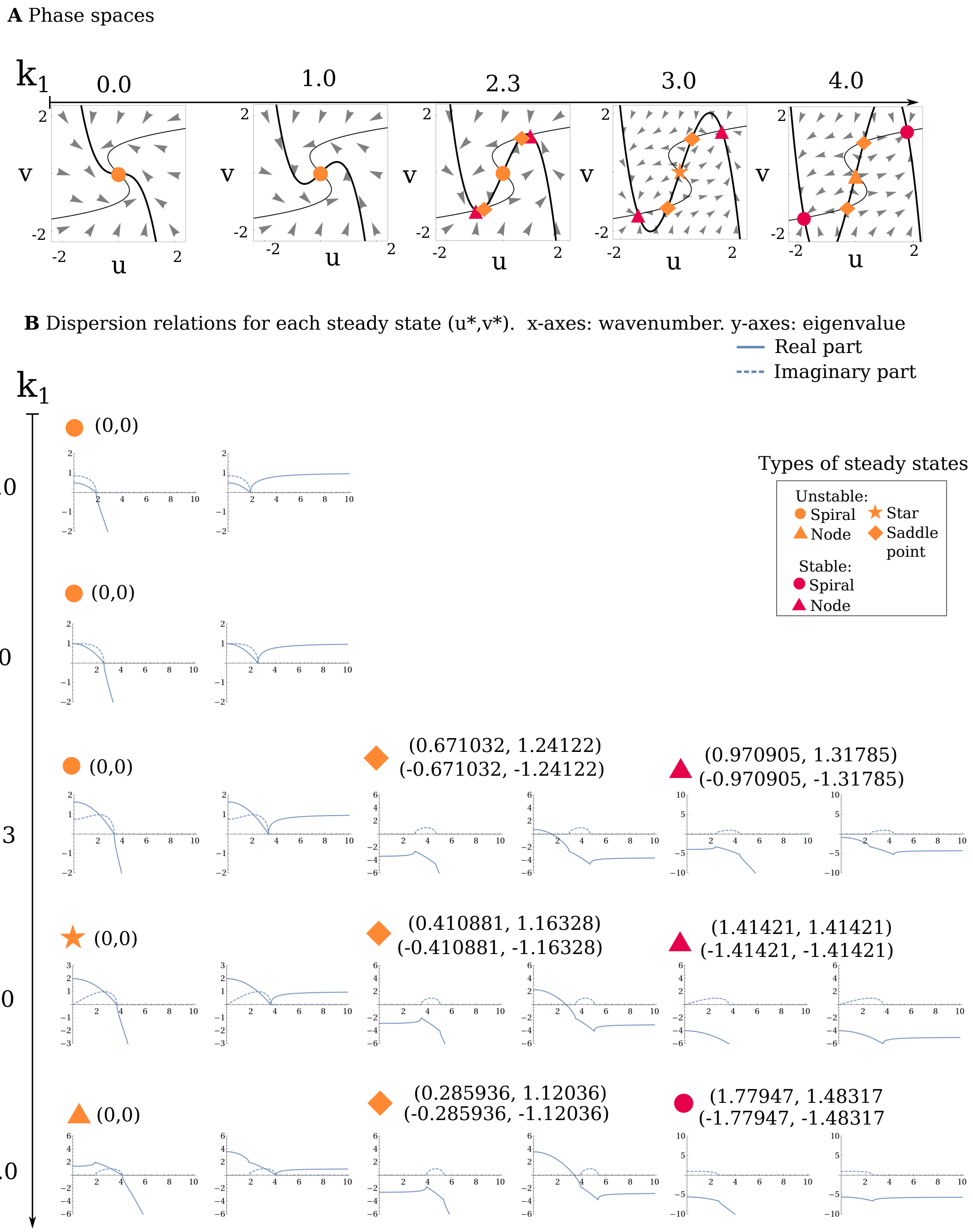


Figure S2: (Caption on next page.)

Figure S2: **Dispersion relation diagrams for the steady states of the example systems in Figure 2E.**
**A)** Phase spaces (as shown in Figure 2F).
**B)** Dispersion relation diagrams show the real (solid line) and imaginary (dashed line) parts of the eigenvalues as a function of the wavenumber (wavenumber = 2Π/wavelength). Where the real part of the eigenvalue is positive, the wavelength (corresponding to the wavenumber) is unstable. If the real part of the eigenvalue is positive for wavenumber=0, the system is unstable without diffusion. If it is negative, the system is stable without diffusion.

### S2 Models that generate rotating wave patterns

**Van der Pol equations** [6]:

$$\frac{dX}{dt} = c(X - X^3/3 + Y) + D_X \nabla^2 X, \tag{S3}$$

$$\frac{dY}{dt} = -X/c. \tag{S4}$$

We have simulated rotating wave patterns with the Van der Pol equations with the parameters: $D_X = 0.3$, dt=0.002, $c = 1$, $L = 100$, with initial noise around $(X, Y) = (0, 0)$ and zero flux boundaries.

**Brusselator equations** [15]:

$$\frac{dX}{dt} = A + X^2Y - (B + 1)X + D_X \nabla^2 X, \tag{S5}$$

$$\frac{dY}{dt} = BX - X^2Y + D_Y \nabla^2 Y. \tag{S6}$$

We have simulated periodic wave patterns containing spirals with the Brusselator equations with the parameters found in the paper by Torabi and Davidsen[20]: $A = 1.9$, $B = 4.8$, $D_X = 1.0$ and $D_Y = 0.7$, $L = 256$. We set the initial values around $(X, Y) = (1.9, 2.52632)$ and used zero flux boundaries.

**The Sevilletor equations**:
Equations (1) and (2) repeated for convenience:

$$\frac{du}{dt} = k_1 u - v - u^3 + D\nabla^2 u, \tag{S7}$$

$$\frac{dv}{dt} = v + u - v^3. \tag{S8}$$

We simulate rotating waves with $k_1 = 1.0$ and periodic wave patterns with $k_1 = 2.3$. $D = 0.3$, $L = 100$. The initial values are around $(u, v) = (0, 0)$ and the system has zero flux boundaries.

**Differences between the Sevilletor model and classic models of the Belousov–Zhabotinsky reaction**
The pattern generated by the Sevilletor model near the bifurcation point ($k_1 = 2.3$) is a new type of spiral patterning behavior that to the best of our knowledge has not been described previously (Figure 2E third row and Movie 1). Although the pattern looks similar to those formed by classic models of the Belousov–Zhabotinsky reaction [22] such as the Brusselator [14] and Oregonator [5], it emerges from a different dynamical behavior. The classic models [14, 5] generate spiral patterns based on a limit cycle around a single unstable steady state (Figure S3), as shown for the rotating pattern presented in this paper with $k_1 = 1$ (Figure 3C). In contrast, the periodic wave patterns with spirals formed by the Sevilletor model with $k_1 = 2.3$ arises from a diffusion-driven excitation of a bistable regime (Figure 3D and Movie 3).

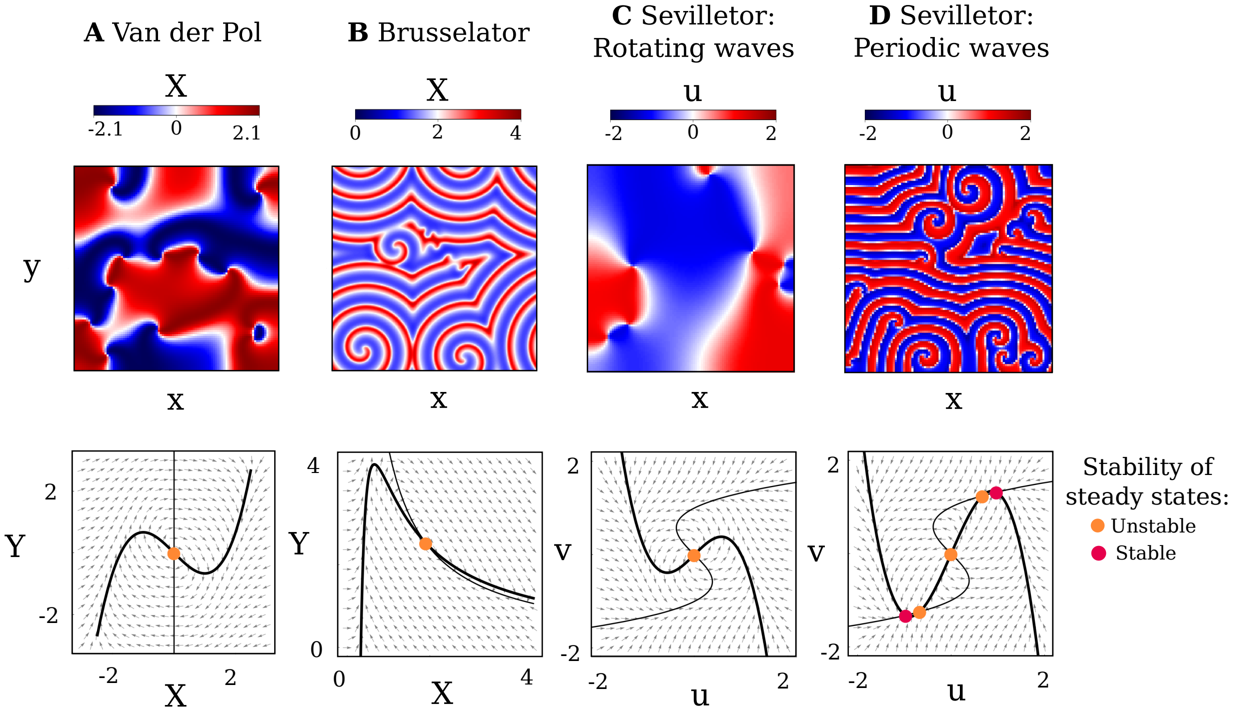


Figure S3: **Other models that produce rotating wave patterns.**
Simulations of rotatory wave formation (top row) and corresponding phase spaces (bottom row). The Van der Pol[6] (A), Brusselator[15, 20] (B) and Sevilletor rotating waves (C) have a single unstable steady state and a limit cycle in the phase space. The Sevilletor periodic waves (D) has five steady states, including two stable ones. However, visually the Van der Pol is similar to (C), and the Brusselator similar to (D).

This difference is also reflected by different patterning dynamics. Our analysis shows that in the Brusselator model, spirals are formed out of local heterogeneities in the initial conditions. Analysis of the phase space shows that these neighboring cells follow a spiral trajectory towards the central steady state, temporarily transforming the unstable steady state of the system into an effectively stable one. Following this trajectory, neighboring cells reach the steady state and eventually synchronize their phases following the original limit cycle. This event is associated with the disappearance of spiral centers. These initial spiral centers are formed independently of boundary conditions and are required produce a stripy phase pattern.

In contrast, with $k_1 = 2.3$, the Sevilletor model generates spirals that appear and disappear dynamically over time while moving in space (Movie 1). Numerical simulations show that these spirals are formed only with zero flux boundary conditions, suggesting that the spiral appearance is due to crashing of phase waves reflected by zero flux boundary conditions. With periodic boundary conditions, the long term behavior of the system is to generate a periodic phase wave pattern of straight stripes (Figure S4).

**FitzHugh-Nagumo model**:
Like the Sevilletor model, another model that was also developed from the Van der Pol equations is the famous FitzHugh–Nagumo model, proposed by FitzHugh in 1961 [6] to study neuronal firing. In this model, an additional excitation term in equation (1) and a negative linear term in equation (2) was added to investigate how a stable system could be excited to become unstable. Unlike the FitzHugh–Nagumo model, the Sevilletor model always has an instability in the steady state $(u^*, v^*) = (0, 0)$ that is driven by the two positive self-enhancing feedbacks of $u$ and $v$. Moreover, while the FitzHugh–Nagumo model can have only up to three steady states, due to the linear shape of the $v$ nullcline, in the Sevilletor model both $v$ and $u$ have nullclines with non-linear shapes that

can intersect on more than three points, which is fundamental to give rise to the excitble bistable behavior.

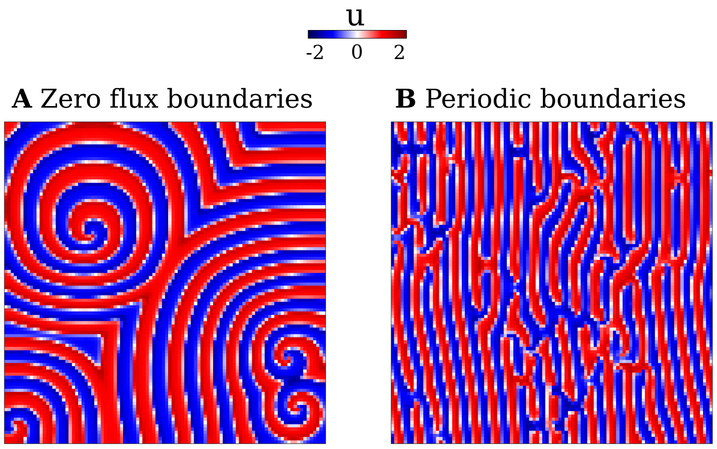


Figure S4: **The Sevilletor periodic wave behavior does not create spiral centers in systems with periodic boundaries.** The Sevilletor with $k_1 = 2.3$ in equations (1) and (2) in systems with **A)** zero flux boundaries and **B)** periodic boundaries.

#### S3 Parameter space that gives rise to periodic wave patterns

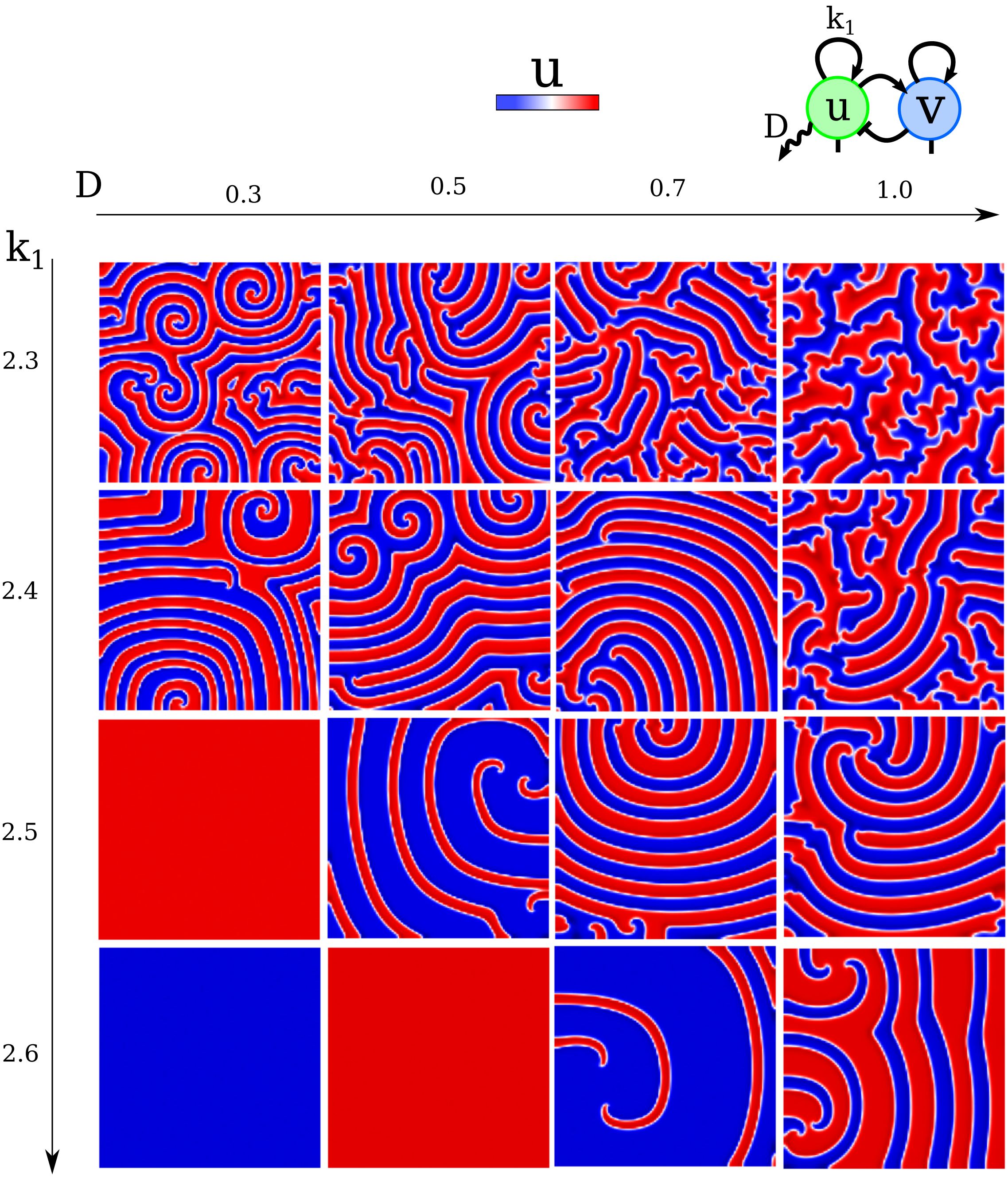


Figure S5: **Parameter values of $k_1$ and $D$ that give rise to periodic wave patterns**. As showed in Figure 3D, periodic wave patterns arise near the bifurcation point between oscillations and bi-stability ($k_1 = 2.3$), where diffusion can destabilize the system to generate a new limit cycle. This behavior is possible for a broad range of values of $k_1 \geq 2.3$ provided that the destabilization by diffusion is strong enough $D \geq 0.3$.

#### S4 Initial values of u and v determine different types of spatial synchronizations

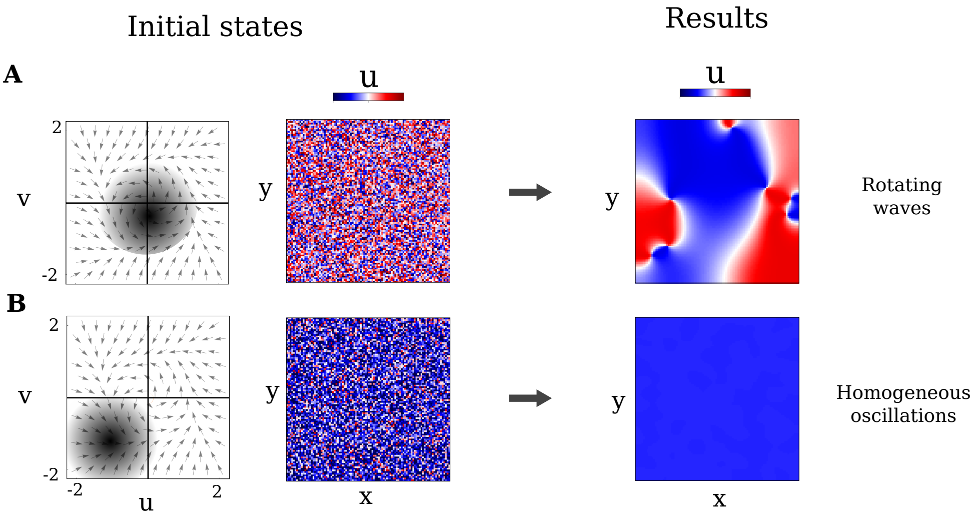


Figure S6: **The type of spatial synchronizations are determined by the initial concentrations.**
The self-organizing patterning formed by the Sevilletor model with $k_1 = 1$ is influenced by the half-planes where initial conditions lie. Left column: initial concentrations illustrated in the phase space (steady states are not shown). Middle column: Corresponding initial conditions for the 2D simulation. Right: Result of the corresponding 2D simulation.
**A)** Uniformly distributed random initial conditions around $(u, v) = (0, 0)$ that lie both in lower and upper half planes give rise to rotating wave patterns.
**B)** Uniformly distributed random initial conditions around $(u, v) = (-1, -1)$ that lie only on the lower half plane give rise to homogeneous oscillations. The same result can be obtained with initial conditions that lie in the upper half plane.

#### S5 Adding a self-regulatory delayed negative feedback of Notch does not change the oscillatory behavior of Wnt and Notch

In classical somitogenesis models oscillations are driven by a cell autonomous transcriptional feedback of Notch mediated by repressors of the Hes/Her family [18, 2]. This transcriptional inhibition can be implemented with a delayed negative feedback that generates sustained oscillations [11] (Figure S7A):

$$\frac{d\texttt{Notch}}{dt} = -\texttt{Notch}_{delay}(\tau - 15) \tag{S9}$$

In the Sevilletor equations, we assume that the additional negative feedback that couples Wnt and Notch complements the core delayed negative feedback of Notch. This additional feedback is implemented by two simple linear interactions with opposite sign and parameters $k_3$ and $k_4$, generating an effective delayed negative feedback that gives rise to out of phase oscillations of Wnt and Notch (Figure S7B).

$$\frac{d\texttt{Wnt}}{dt} = \texttt{Wnt} - k_3\texttt{Notch} - \texttt{Wnt}^3 + D\nabla^2\texttt{Wnt} \tag{S10}$$

$$\frac{d\texttt{Notch}}{dt} = \texttt{Notch} + k_4\texttt{Wnt} - \texttt{Notch}^3 - \texttt{Notch}_{delay}(\tau - 25) \tag{S11}$$

The delayed negative feedback that couples Wnt and Notch is sufficient to drive oscillations (Figure S7C). Together with the self-enhancement, this feedback generates self-organizing patterns due to synchronizations mediated by diffusion, see Figure 2 and Figure 3. To maintain the model complexity to a minimum, we consider this reduced self-organizing model without explicitly including the direct delayed negative feedback of Notch.

$$\frac{d\texttt{Wnt}}{dt} = \texttt{Wnt} - k_3\texttt{Notch} - \texttt{Wnt}^3 + D\nabla^2\texttt{Wnt} \tag{S12}$$

$$\frac{d\texttt{Notch}}{dt} = \texttt{Notch} + k_4\texttt{Wnt} - \texttt{Notch}^3 \tag{S13}$$

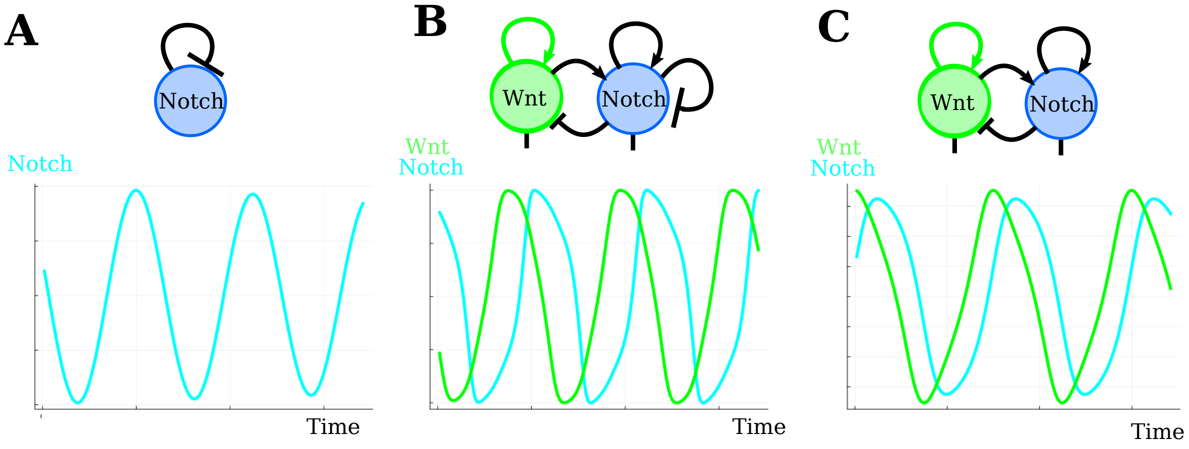


Figure S7: **The Sevilletor includes a delayed negative feedback between Wnt and Notch, complementing the direct delayed negative feedback of Notch**.
**A)** A delayed negative self-regulation of Notch gives rise to oscillations.
**B)** Including an additional delayed negative feedback that couples Wnt and Notch as in Sevilletor network does not affect the oscillatory behavior of the system ($k_3 = k_4 = 1$).
**C)** The delayed negative feedback that couples Notch and Wnt is sufficient to drive out of phase oscillations of Wnt and Notch ($k_3 = k_4 = 1$).

### S6 The original PORD model by Cotterell et al. [3]

The PORD model by Cotterell et al. [3] freezes cells in opposite phases as observed in the lateral inhibition patterning regime of the Sevilletor model, as shown by the two cell simulations in Figure S8A. The cell starting in (0,0) makes one single loop around the steady state, before freezing at the bottom left corner while the other cell freezes at the top right corner. This is a non-symmetric phase space, while in our PORD implementation (Figure S8B) the phase space and the path of cells are symmetric, starting from opposite sides of the steady state. This difference, together with use of a heavy side function to prevent negative values, may explain why the periodic peaks formed by the original PORD model are separated by several cells, while in our model and in other implementations of the PORD hypothesis [9] they are separated by only one cell. Despite this difference, all PORD implementations exhibit the same underlying behavior based on the freezing of oscillations in opposite phase due to diffusion between neighboring cells.

The equations of the original PORD model by Cotterell et al.[3] are:

$$\frac{dA}{dt} = \frac{\Phi\big(k_1 A + k_2 R + F + \beta\big)}{1 + k_1 A + k_2 R + F + \beta} - \mu A, \tag{S14}$$

$$\frac{dR}{dt} = \frac{k_3 A}{1 + k_3 A} + D_R \nabla^2 R - \mu R. \tag{S15}$$

Here $\Phi(x) = x \cdot H(x)$, where $H(x)$ is a Heavyside function: $H(x) = 1$ for $x > 0$ and $H(x) = 0$ for $x \leq 0$. The parameters used in Figure S8 are the following: $k_1 = 1.56$, $k_2 = -2.28$, $k_3 = 0.099$, $\beta = 0.5$, $\mu = 0.05$, $F = 1$. $D_R = 0.008$.

The Sevilletor equations for the lateral inhibition behavior used to make an implementation of the PORD model are (equations (1) and (2) repeated for convenience):

$$\frac{du}{dt} = k_1 u - k_3 v - u^3 + D\nabla^2 u, \tag{S16}$$

$$\frac{dv}{dt} = k_2 v + k_4 u - v^3. \tag{S17}$$

With $k_1 = 0.0, k_2 = 1$ and $D = 1.0$. For the example shown in Figure S8 we set the parameters of the frequency gradient $k_3 = k_4 = 1.0$.

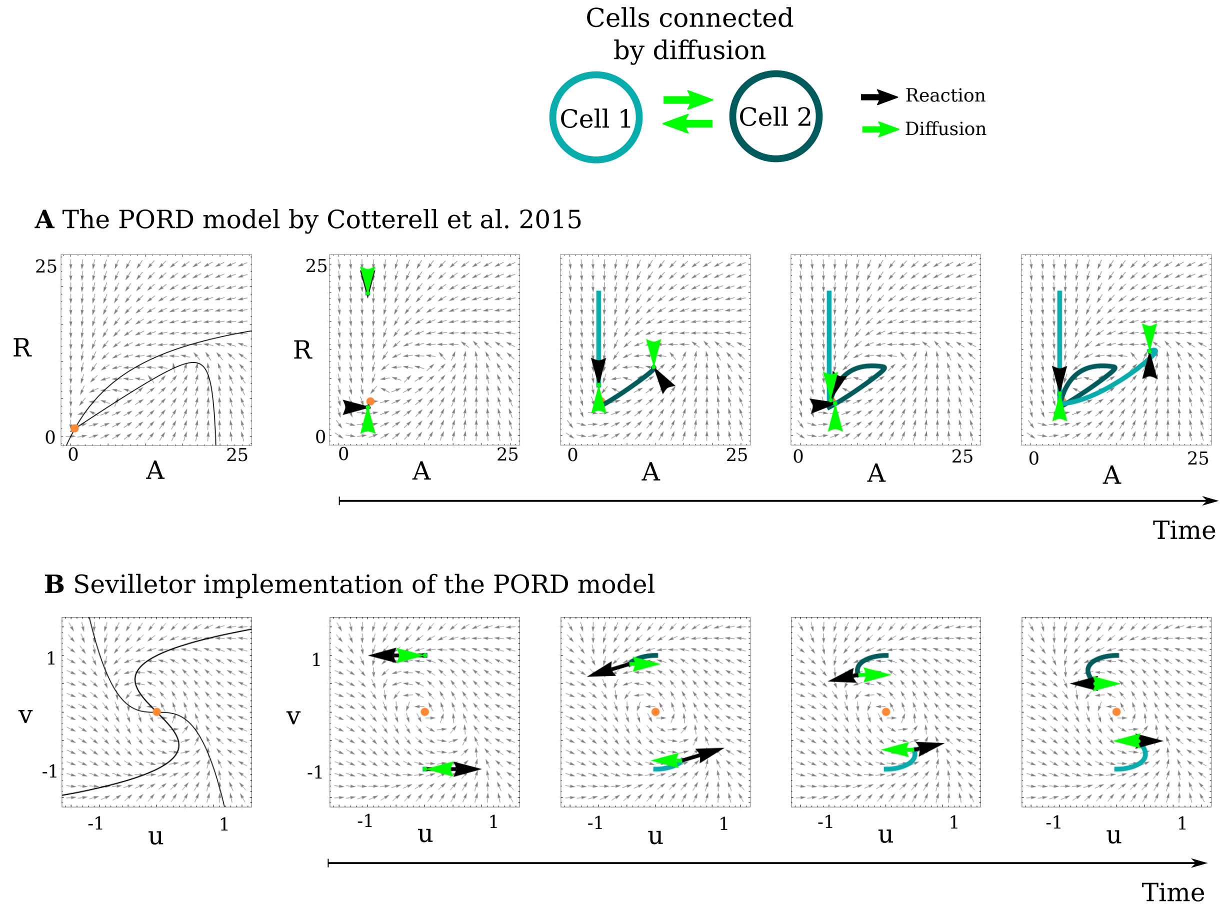


Figure S8: **The original PORD Model by Cotterell et al.[3] and the Sevilletor implementation.**
**A)** The original PORD model by Cotterell et al. [3] and **B)** our implementation in the Sevilletor framework with $k_1 = 0$, $k_2 = 1$, $k_3 = 1$ and $k_4 = 1$. Left: Phase spaces with nullclines (black lines) and unstable steady states (orange). Right: The time series show the path of 2 cells with initial values: A) $(A, R) = (0, 20)$ and $(0, 0)$, B) $(u, v) = (0, 1)$ and $(0, -1)$. In both cases the cells freeze out of phase when reaching a point where changes promoted by reactions (black arrows) are counterbalanced by diffusion (green arrows), freezing cells in opposite state in both models.

The frequency gradient is implemented in different ways in the original and the Sevilletor PORD model. In the original PORD model it is a gradient of the strength of the parameter $F$ which enhances $A$, while in the Sevilletor implementation it is a graded modulation `PG` of the strength of the negative feedback between the two reactants ($k_3$ and $k_4$).

Both the original PORD equations and the Sevilletor implementation are fragile to noise, as shown in Figure S9.

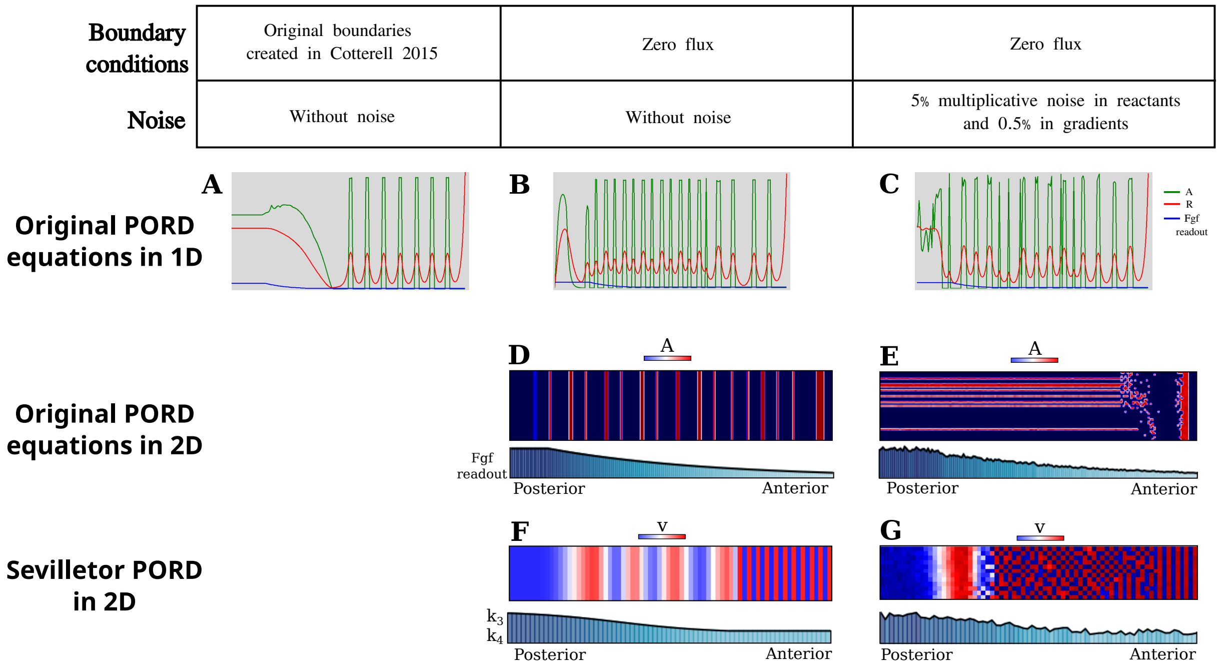


Figure S9: **Noise in the PORD model creates a disrupted pattern.**
**A-C)** The original PORD model by Cotterell et al. [3] in 1D (equations (S14) and (S15)). The parameters used are the following: $k_1 = 1.56$, $k_2 = -2.28$, $k_3 = 0.099$, $\beta = 0.5$, $\mu = 0.05$, $F$ is a gradient from 1 to 0 and $D_R = 1.0$. An initial peak of $R$ is placed at the most anterior line.
**A)** A regular striped pattern is formed with the original boundary conditions created by Cotterell et al. [3] and without noise.
**B)** A striped pattern with variations in somite width is formed with zero flux boundary conditions and without noise.
**C)** An irregular striped pattern is formed with zero flux boundary conditions and with 5% multiplicative noise in the concentrations of A and R and 0.5% multiplicative noise in the gradient of Fgf readout ($F$).
**D-E)** The original PORD equations by Cotterell et al. [3] in 2D (equations (S14) and (S15)).
**D)** A striped pattern with variations in somite width is formed with zero flux boundary conditions and without noise.
**E)** A disrupted pattern is formed with zero flux boundary conditions and with 5% multiplicative noise in the concentrations of A and R and 0.5% multiplicative noise in the gradient of Fgf readout.
**F-G)** The PORD model in the Sevilletor framework in 2D.
**F)** A regular striped pattern is formed with zero flux boundary conditions and without noise.
**G)** A disrupted pattern is formed with zero flux boundary conditions and with 5% multiplicative noise in the concentrations of $u$ and $v$ and 0.5% multiplicative noise in the gradient of $k_3$ and $k_4$.

### S7 The CWS model is robust to noise

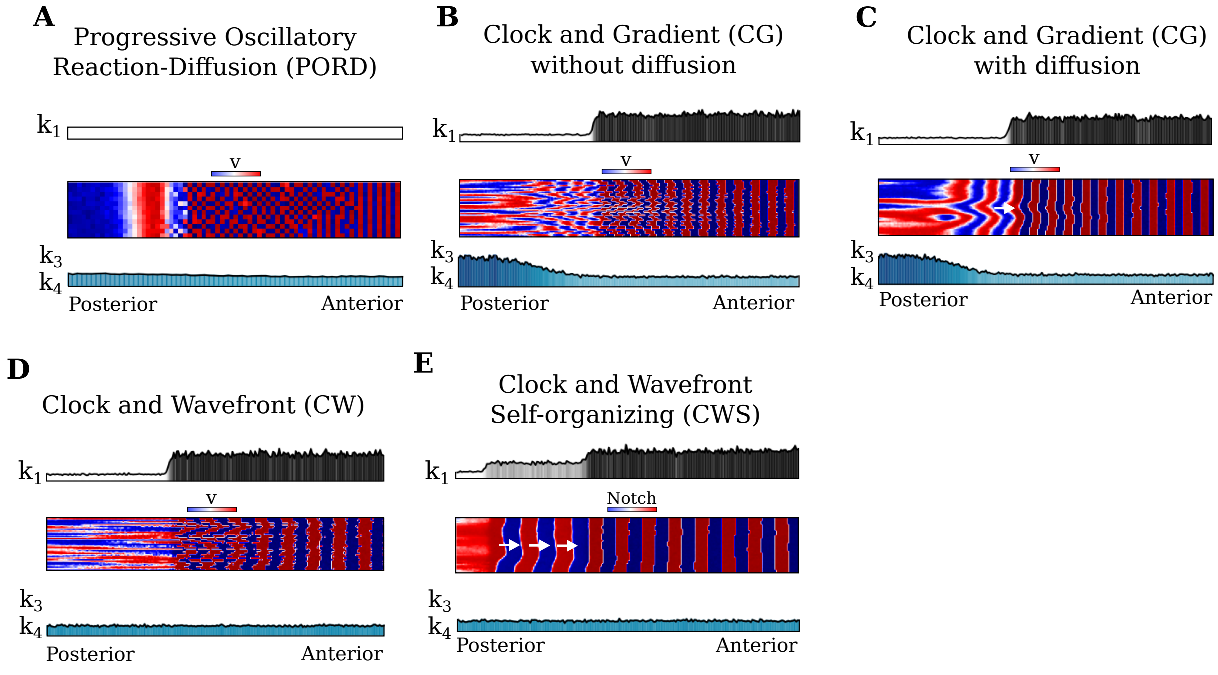


Figure S10: **Models of somitogenesis with multiplicative noise throughout the simulation**. $v$ signaling patterns in the CW, CG, PORD and CWS model with noise added to the values of $u$ and $v$, frequency gradients (values of $k_3$ and $k_4$) and the parameter that defines regions with different dynamical regimes (values of $k_1$) at every time when the tail grows from a Gaussian distribution. 5% noise is added to $u$ and $v$ and 0.5% noise is added to $k_1, k_3$ and $k_4$.
**A)** In PORD models, low amounts of noise disrupt the somite pattern formation and generates a chessboard pattern (Movie 6). In this case no noise is added to $k_1$ as it is set to 0 everywhere. The same result is found for the original PORD equations in Cotterell et al. [3], as shown in Figure S9.
**B-C)** In the CG model, the accumulated noise in the frequency gradient eventually disrupts the somite pattern (Movie 7). Diffusion in the CG model allows for self-organizing coordination between cells, prolonging the time it takes for the pattern to be disrupted by noise compared with the example shown without noise (B).
**D)** The CW model shows a low robustness to noise.
**E)** The CWS model generates normal somite patterns with a greater robustness to noise because the phase wave propagation is independent from global frequency gradients (Movie 8).

### S8 Dissection of the tailbud in the CWS model

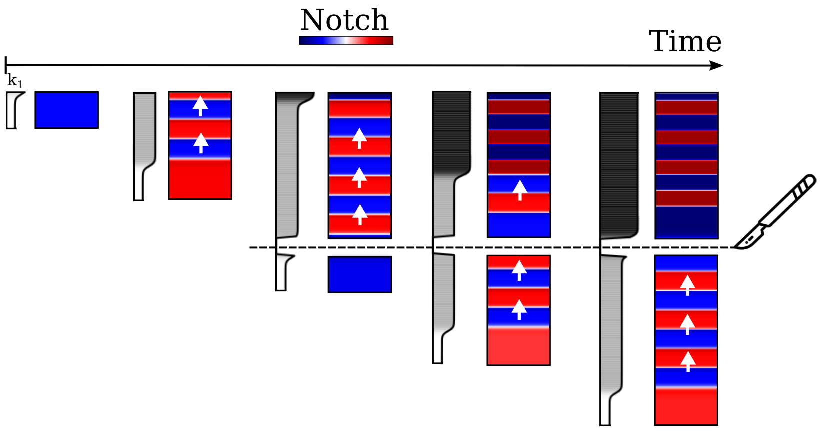


Figure S11: **Virtual dissection of the tailbud in the CWS model (Movie 10).** Time series showing the effect of tailbud removal in a simulation using the Clock and Wavefront Self-Organizing model. Following the separation of the posterior region, the tailbud continues to grow and to generate new phase waves that travel anteriorly. In the anterior dissected part, preexisting phase waves persist and propagate anteriorly, driven by the excitable behavior in the intermediate region (light gray section), until they stop when entering the bistable region (dark gray section). These findings align with experimental observations reported in [23]. Importantly, in the anterior dissected part, the propagation of straight waves without rotational movements occurs independently of the homogeneous oscillations at the posterior tailbud tip.

#### S9 Adding diffusion of Notch to the CWS model

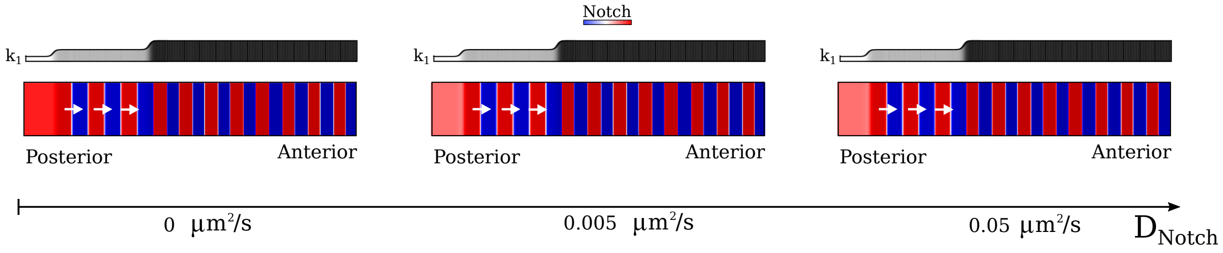


Figure S12: **Adding diffusion of Notch to the CWS model does not affect the overall dynamics**. Interpreting $u$ as Wnt and $v$ is Notch in the CWS model, the diffusion constant of Wnt is estimated in Section S17 to be $\mathtt{D_{Wnt}} = 0.048\ \mu\text{m}^2/\text{sec}$. Including diffusion of both reactants does not change the phase waves and somite pattern formation in the CWS model.

#### S10 CWS model with thinning of the waves along the anterior-posterior axis

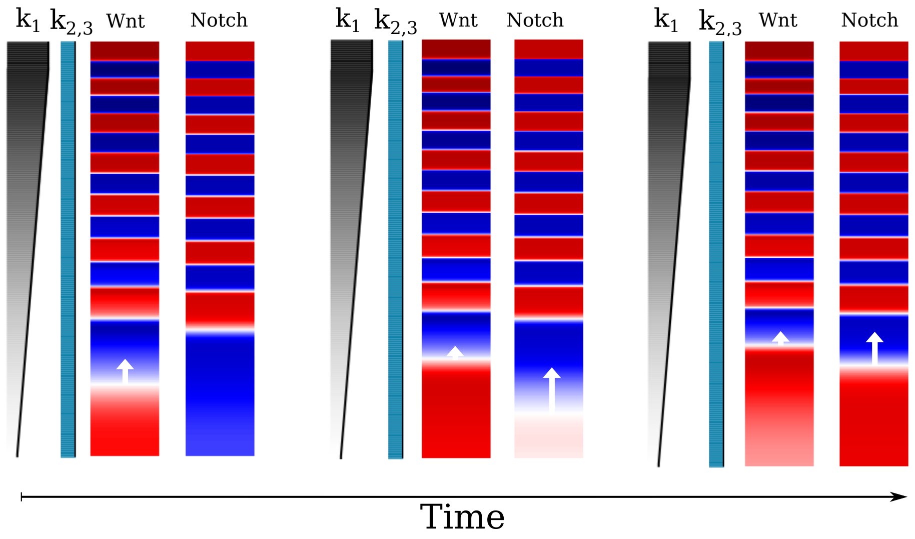


Figure S13: **CWS model with thinning of the waves along the anterior-posterior axis**. Timeline of the CWS with a graded change in $k_1$ creates waves with long wavelengths at the posterior tip that become narrower as they approach the anterior side (Movie 9). Wnt initiates the wave, which slows down as it moves anteriorly, when the wave of Notch rapidly catches up.

### S11 The CWS model recapitulates the rotating wave patterns observed in mixed explant experiments

Ex-vivo cultures of mouse presomitic mesoderm (PSM) offer valuable insights to explore the coordination of oscillations during somitogenesis. An explant is formed by dissecting a portion of the embryonic tail, creating a circular quasi-monolayer culture in-vitro [21] (Figure 5F). In this section, we present virtual explant simulations that mirror experimental conditions, projecting the entire posterior tail into a circular domain, with the posterior tailbud at the center and the intermediate part at the outer edge (Figure 5E,G).

Explants have been generated using alternative protocols that dissected either the middle [10] or entire posterior part of the tail [7]. In both cases, sequential signaling waves resembling somitogenesis propagated from the inner to the outer part of the explants. However, in the former case, oscillations ceased after a few cycles concomitantly with segment boundary formation [10], while in the second case, they persisted for up to twenty cycles over two days [7]. The mechanisms underlying wave self-organization in explants remain unclear. In the former case, evidence suggests that cells may re-establish posterior signaling gradients [10, 21], while in the latter, homogeneous distribution of signals and their targets indicates no contribution to spatial modulation [7]. Discrepancies may arise from specific culture conditions, like removing the ectoderm and supplementing media with activators and inhibitors of BMP, FGF, WNT, RA, and ROCK, as performed in the second case.

Since the properties inherited or re-established by cells in the tail are unknown, we conducted virtual explant simulations with decreasing degrees of information (Figure 5G). In all simulations, cells inherit the dynamical behavior that is characteristic of each model ($k_1$). However, they can retain the positional information provided by gradients ($k_3$ and $k_4$) and phase values ($u$ and $v$ concentration), only the phase values, or no information (Figure 5E,G). We generate explants of the PORD, CG and CWS models. To introduce a minimal degree of cell mixing similar to real explants, we assumed that cells switch positions randomly with their neighbors (Figure 5G). In addition, we vary the size of the posterior region that is projected from the tail (Figure S14), we included explants with only the middle portion of the tail (Figure 5L-N), and we simulate explants where all cells are mixed together (Figure S15).

#### S12 Varying the position of the cut in explants

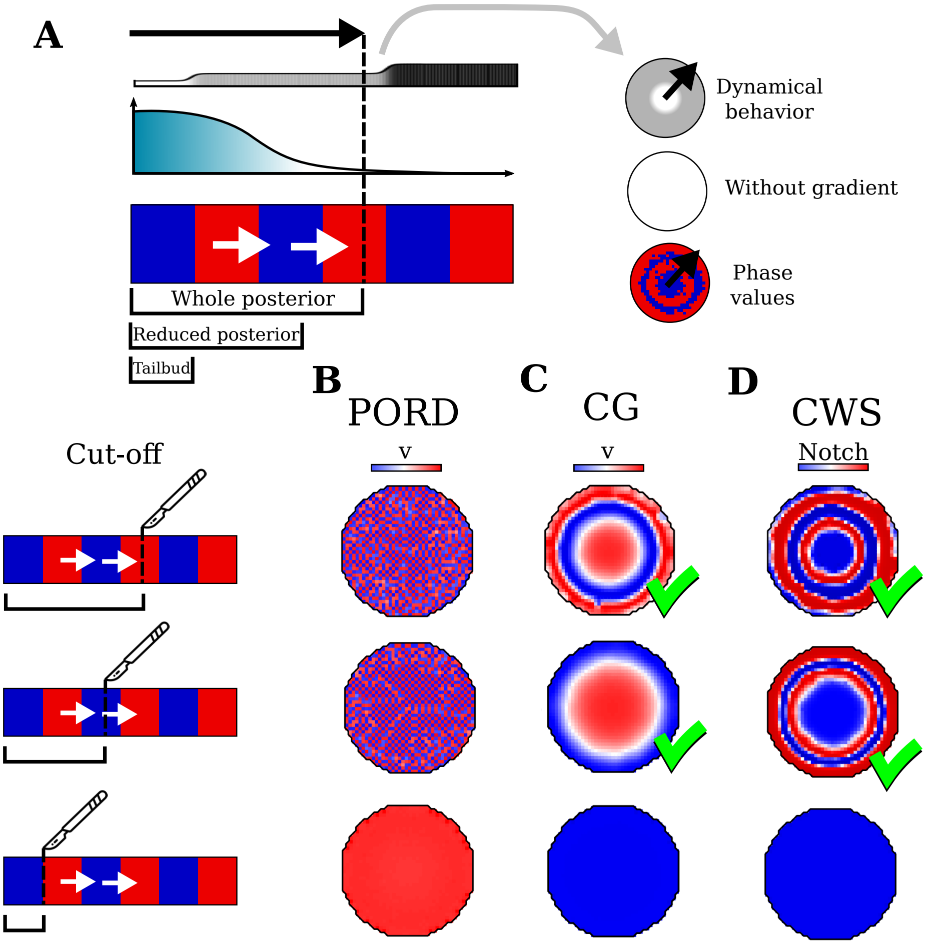


Figure S14: **Varying the position of the cut in explants.**
**A)** A drawing showing different virtual explants obtained by cutting the tail at different positions in the PORD, CG and CWS model. Explants retain dynamical behaviors ($k_1$) and phase values ($u$ and $v$ concentrations).
**B-D)** The different explants generate the same patterning behaviors as shown in Figure 5, except the case when only the posterior tailbud is cut (lower row) where all the models generate homogeneous oscillations.

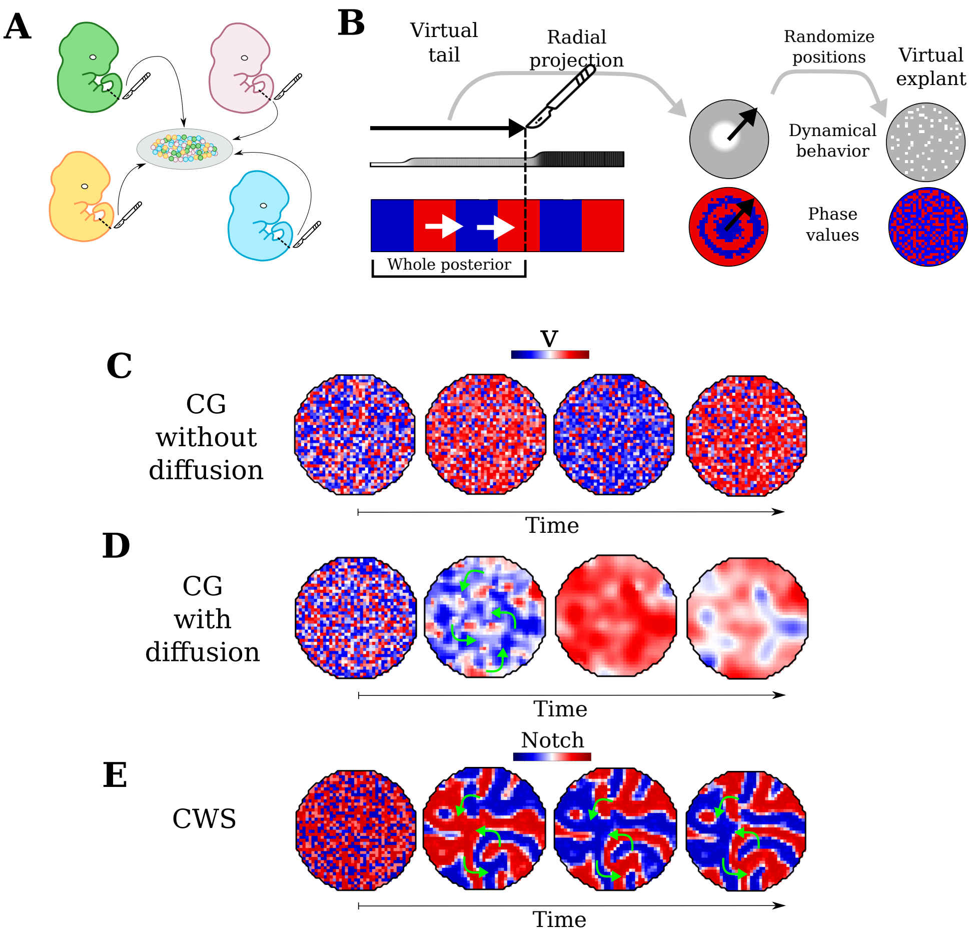


Figure S15: **The CWS model recapitulates the self-organizing behavior observed in mixed explant experiments (Movie 13).**

**A)** Illustration of experiments where explants are obtained by mixing cells from different tails.

**B)** Virtual mixed explants of the CG and CWS model are performed by projecting a portion of the tail radially and by randomizing the positions of the cells to mimic experiments where cells are mixed from different tails. The randomized cells in the explant retain the dynamical behavior and phase values ($u$ and $v$) possessed in the tail.

**C)** Mixed explants in a cell-autonomous CG model without diffusion generate salt and pepper patterns where each cell continue individual oscillations depending on the initial phase value inherited from the tail.

**D)** Mixed explants in CG model with diffusion. Local synchronizations between cells promote rotating patterns (green arrows) (Movie 13).

**E)** Mixed explants of the CWS model generate periodic phase waves with a sustained rotating pattern (green arrows) (Movie 13).

#### S13 Ablating the center of explants from the Clock and Wavefront Self-Organizing model

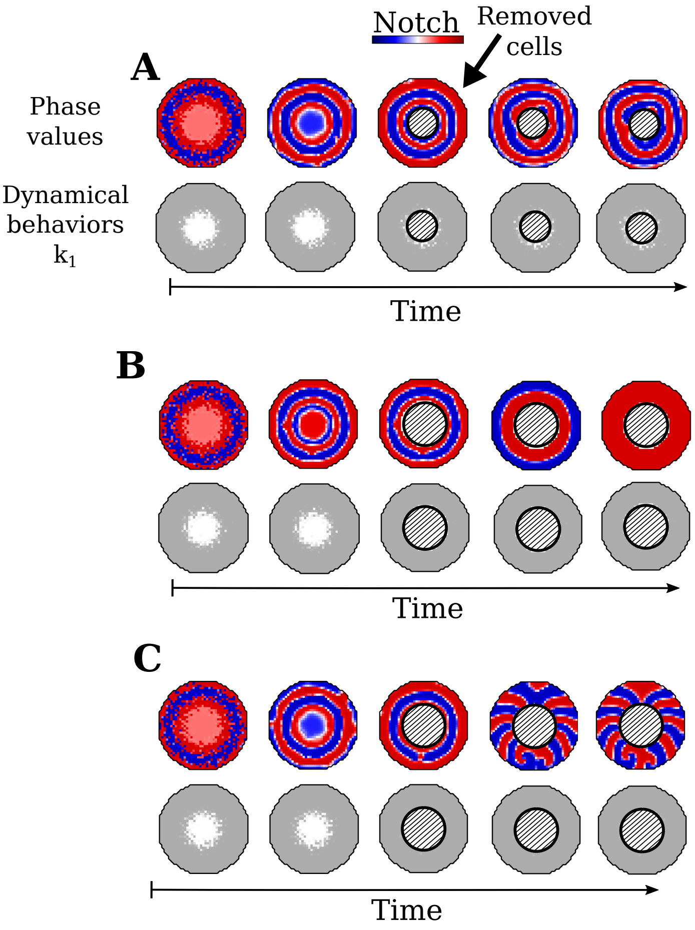


Figure S16: **Phase waves still propagate after ablating the center of explants derived from the Clock and Wavefront Self-Organizing model**.
Time series of Notch ($v$) in explants of the Clock and Wavefront Self-Organizing model after ablation of the center cell population.
**A)** When the vast majority of cells in the center is ablated (white cells in the bottom row), CWS explants keep propagating target patterns. The presence of a few residual central cells (residual white cells the bottom row) is sufficient to always sustain target patterns.
**B-C)** If a larger central portion is ablated in the CWS explants, the circular target patterns keep propagating initially, but depending on phase values they can either dissipate into homogeneous patterns (B) or evolve into periodic phase wave patterns that exhibit rotations (C). This is similar to the behavior observed in explants obtained solely with cells derived from the middle part of the tail (Figure 5N). In all cases, the cells in the outer layer form phase waves in a self-organizing manner independently of global frequency gradients.

### S14 Possible outcomes of middle tail explants in CWS model

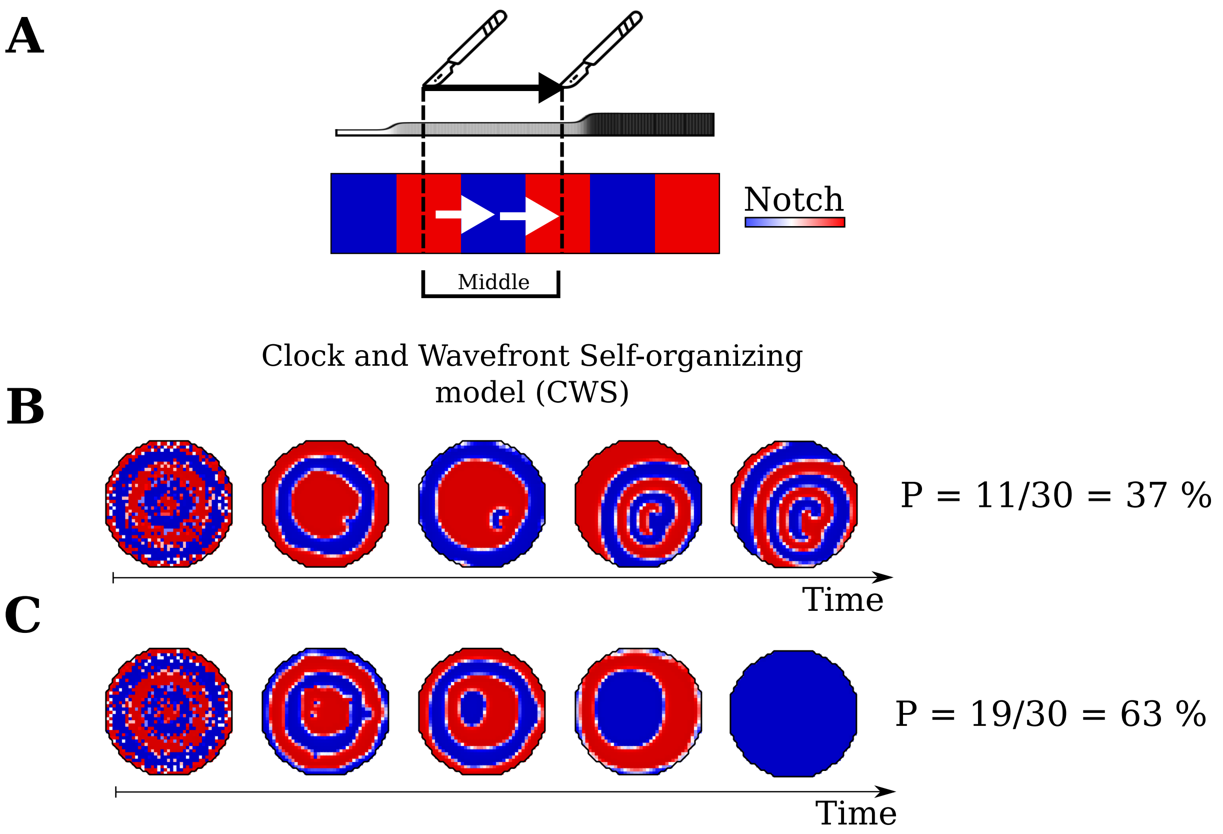


Figure S17: **Variability in virtual middle explants of the Clock and Wavefront Self-Organizing model.**
**A**) Illustration of the part of the tail used to create an explant of the Clock and Wavefront Self-Organizing model that only includes cells with excitable behavior in the middle of the tail.
**B-C**) Summary of 30 simulations of the CWS model performed where the variability across simulations depends on the stochastic shuffling between cells performed in the making of the explants. Explants obtained from the middle section of the tail (without the tip) give rise to two possible outcomes: circular wave patterns with a single center made from two spirals with opposite rotation directions (Movie 11) in 37% of simulations (11/30), or a set of concentric waves that dissipates into a homogeneous pattern in 63% of the cases (19/30). This shows that when the pacemaker population from the center of the explant is not included, self-organizing target patterns can arise solely from the initial phase values inherited from the tail, but that their evolution depends on the specific initial distribution of phase values.

235 ## S15 Reversed negative feedback between Wnt and Notch in the
236 CWS

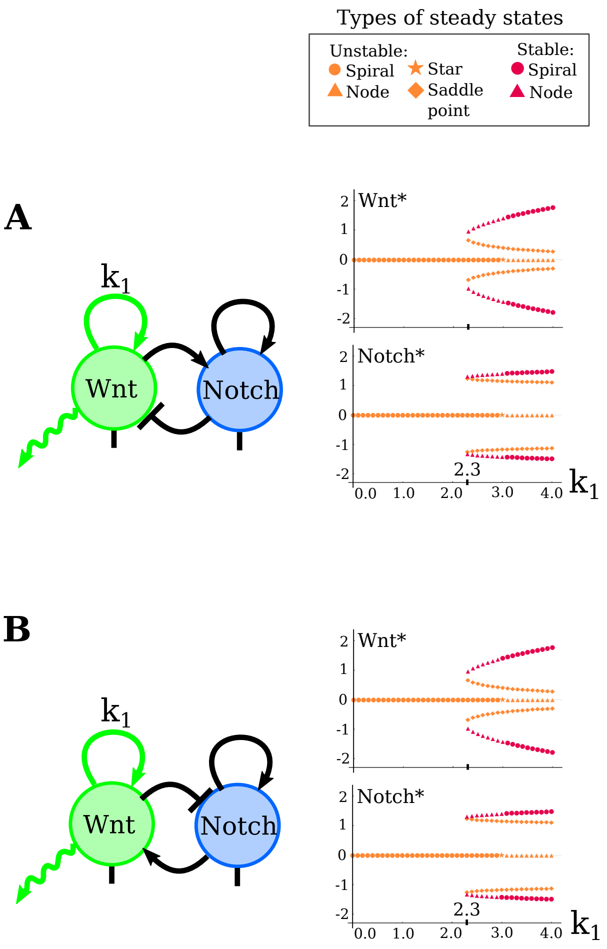


Figure S18: **Alternative implementation of the CWS model where the signs of the negative feedback loop between Wnt and Notch are inverted.**
First column: Network diagrams with a standard (A) or reversed (B) feedback between Wnt ($u$) and Notch ($v$). Second column: The bifurcation diagrams of Wnt and Notch associated with the parameter $k_1$. Third column: somitogenesis simulations of the CWS model with a representation of the modulation of $k_1$ in the model (top line) and Wnt and Notch signaling patterns (second and third lines). Check marks show whether the different implementations of the model can reproduce the Wnt and Notch relative phases observed at the posterior, middle and anterior part of the tail in experiments [1, 17].
**A**) The standard implementation of the model presented in Figure 4 with a positive interaction from Wnt to Notch and an inhibition from Notch to Wnt. The CWS model can generate realistic patterns with the correct relative phase between Notch and Wnt at the posterior, middle and anterior part of the tail (three green marks), in agreement with experimental data [1, 17].
**B**) When the incoherent negative feedback between Notch and Wnt is inverted, the CWS model cannot reproduce the relative phase change between Notch and Wnt observed in experiments.

### S16 Wnt patterns in models of virtual tails and explants

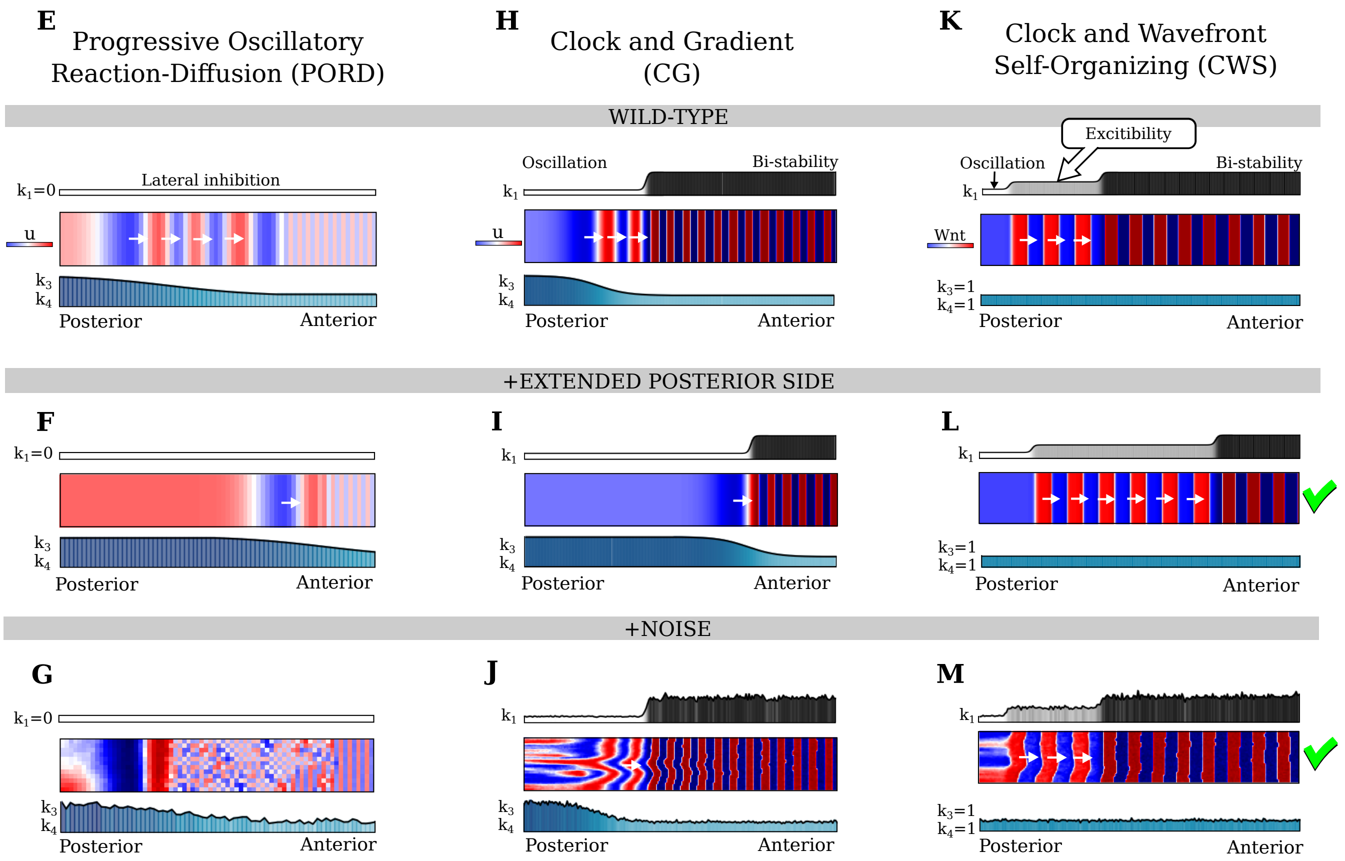


Figure S19: **The models of somitogenesis in virtual tails of Figure 4 shown for** $u$**.** See the Caption of Figure 4.

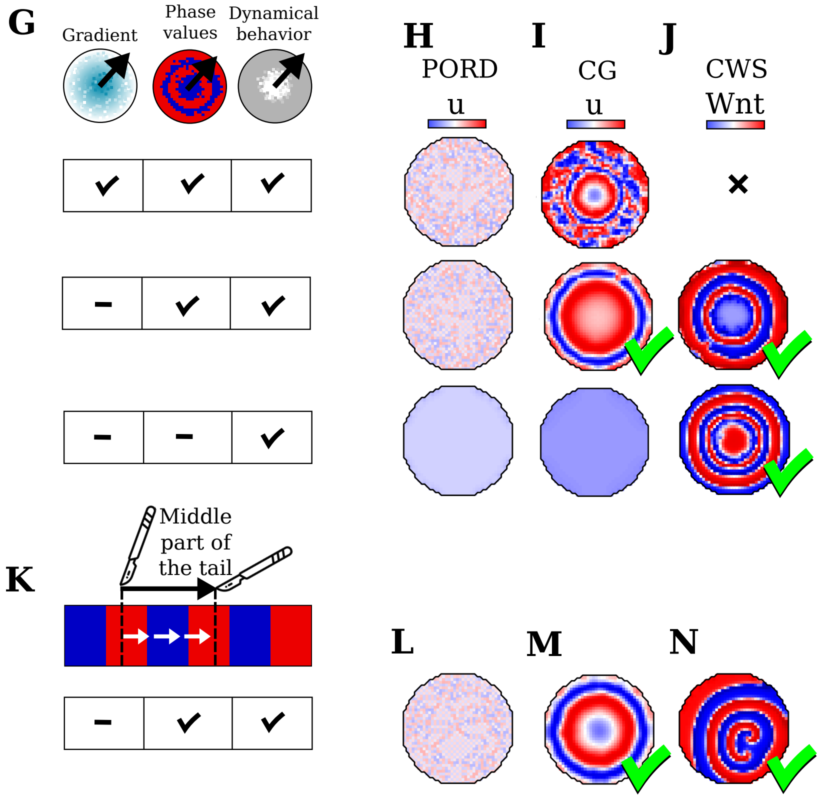


Figure S20: **The models of wave formation in virtual explants in Figure 5 shown for $u$.** See the Caption of Figure 5.

#### S17 Units in the CWS model

The corresponding physical units of space and time of the somitogenesis simulation in the Clock and Wavefront Self-organizing model can be estimated in the following way. The average width of a mouse somite is about 200 $\mu$m [19], and a mouse embryonic stem cell has a diameter of approximately 7-17 $\mu$m[13], giving the estimate that a somite is about 20 cells wide. The width of a somite in the simulation is a similar number of about 17 cells. Therefore, in the simulation the diameter of a cell is 1 space unit `dx`; 1 `dx` $\sim$ 12 $\mu$m.

A pre-pattern of a somite is formed every 2 to 3 hours in mice [17, 12], so 8 somites would be formed after about 16-24 h. The CWS can form 8 somites in 607500 time steps of duration 0.002`dt` with appropriately scaled variables and without normalizing all reaction variables with $k_2$. Setting `dt`=1min, the estimated duration of the simulation is `T` = 0.002 `dt` * 607500 time steps = 20.25 h.

A verification of the estimate of the time and space relation with experimental measurements is found by calculating the corresponding diffusion constant of Wnt $\mathtt{D_{Wnt}}$ =0.02 $\mathtt{dx}^2/\mathtt{dt}$ = 0.048 $\mu$m$^2$/sec, which is in the same order of magnitude as the diffusion constant measured of Wg in Drosophila, the equivalent to Wnt in mice, which is 0.05 $\mu$m$^2$/s [8].

### S18 Summary of Figure 4

Table 1: Summary of Figure 4.

| | Phase waves | | Arrest | |
|---|---|---|---|---|
| | Global signals | Self-organizing | Global signals | Self-organizing |
| CW | - | - | x | |
| PORD | x | | | x |
| CG | x | | x | |
| CWS | | x | x | |

#### S19 Captions of Movies

Movies of two-dimensional square simulations of the Sevilletor model, two-cell simulations in phase space and somitogenesis simulations of the tail and tail explants for the PORD, CG and CWS model. The number of time steps in each movie varies to illustrate the dynamics of the patterning processes.

**Movie 1: Self-organizing patterning behaviors of the Sevilletor model**
From left to right, two dimensional simulations of the Sevilletor model started with noisy initial concentrations for increasing values of $k_1$, as shown in Figure 2E. With $k_1 = 0.0$ the model forms a lateral inhibition chessboard pattern, with $k_1 = 1.0$ rotating waves, with $k_1 = 2.3$ periodic wave patterns with spiral centers, with $k_1 = 3.0$ a homogeneous pattern and with $k_1 = 4.0$ bi-stable salt and pepper pattern.

**Movie 2: Phase space trajectories of two cells in lateral inhibition patterning**
Two-cell simulations in phase space for the lateral inhibition regime with $k_1 = 0.0$ in the absence or presence of diffusion, as showed in Figure 3B. The trajectories of the two cells with initial values $(u, v) = (\pm 0.1, 0.0)$ are shown in teal colored lines, the gray point shows the central unstable steady state. Without diffusion (graph on the left) the two cells oscillate by following a limit cycle around the central steady state. In the presence of diffusion (graph on the right) the two cells stop at opposite phase where the changes promoted by reaction (black arrows) are balanced by the changes promoted by diffusion (green arrows). This behavior generates chessboard patterns as shown in Figure 2E.

**Movie 3: Phase space trajectories of two cells with periodic wave excitable patterning behavior**
Two-cell simulations in phase space for the periodic wave regime with $k_1 = 2.3$ in the absence or presence of diffusion, as showed in Figure 3D. The trajectories of the two cells with initial values $(u, v) = (\pm 0.1, 0.0)$ are shown in teal colored lines, the gray points are unstable steady states and white points are stable steady states. Without diffusion (graph on the left) each cell is pushed by reaction (black arrows) into one of the stable steady states. In the presence of diffusion (graph on the right) the influence of reaction is counterbalanced by diffusion (black and green arrows) that pushes each cell onto the trajectory of the other unstable gray point creating coordinated loops. This behavior generates periodic wave patterns with spiral centers from noise as shown in Figure 2E.

**Movie 4: Changing the patterning dynamics by varying the strength of $k_1$**
The behavior of the Sevilletor model can be easily modulated to transition between patterning regimes over time by varying the parameter $k_1$ in all cells. In this example, the model transitions from lateral inhibition pattern, to rotating waves, to periodic waves with spiral centers, to a bi-stable frozen spiral pattern, and back to lateral inhibition pattern: $k_1 = 0.0 \rightarrow 1.0 \rightarrow 2.3 \rightarrow 4.0 \rightarrow 0.0$.

**Movie 5: $k_1$ and $k_2$ determines the self-organizing patterning**
Square two-dimensional simulation with noisy initial conditions and linear modulations of the parameters $k_1$ and $k_2$ along the y- and x-axis, as showed in Figure 3H. The two parameters increase linearly from 0 to 4 promoting transitions between lateral inhibition pattern, rotating waves, periodic wave patterns, homogeneous patterns and bi-stable frozen patterns.

**Movie 6: Sevilletor implementation of the PORD model**
2D simulation of the PORD model implemented in the Sevilletor framework, showed in Figure 4E-G. Left: wild type, middle: posterior expanded mutant, right: with multiplicative noise in $u$, $v$, $k_3$ and $k_4$. $u$ and $v$ signaling are shown side-by-side. All cells have $k_1 = 0.0$ and the simulations start with high signaling at the top boundary (anterior region). The virtual tail grows with a constant speed by addition of a line of cells at the bottom (posterior). A gradient of $k_3$ and $k_4$

changes the frequency along the y-axis to create phase waves. The model generates a periodic somite pattern due to a rely mechanisms based on lateral inhibition. The PORD model is not able to create more phase waves in expanded posterior mutants and it is fragile to noise (Figure S9).

**Movie 7: Sevilletor implementation of the Clock and Gradient (CG) model**
2D simulation of the cell-autonomous Clock and Gradient model implemented in the Sevilletor framework, showed in Figure 4H-J. Left: wild type, middle: posterior expanded mutant, right: with multiplicative noise in $u$, $v$, $k_1$, $k_3$ and $k_4$. $u$ and $v$ signaling are shown side-by-side. The virtual tail grows with a constant speed by addition of a line of cells at the bottom (posterior). The posterior side has $k_1 = 1$ that promotes out of phase oscillations of the two reactants. The anterior side has a bistable behavior that promotes a progressive freezing of oscillations, which generates a periodic somite pattern with $k_1 = 4$. Phase waves are created with a gradient of $k_3$ and $k_4$ that controls the frequency of oscillations. This model cannot make more phase waves in an expanded posterior mutant and noise in the frequency gradient creates a disruption of the phase waves after the formation of several somites.

**Movie 8: The Clock and Wavefront Self-Organizing (CWS) model recapitulates somitogenesis**
Somitogenesis simulations of the novel Clock and Wavefront Self-Organizing model implemented within the Sevilletor framework, showed in Figure 4K-M. Left: wild type, middle: posterior expanded mutant, right: with multiplicative noise in Wnt ($u$), Notch ($v$), $k_1$, $k_3$ and $k_4$. Wnt and Notch signaling are shown side-by-side. The virtual tail grows with a constant speed by addition of a line of cells at the bottom (posterior). The posterior side has $k_1 = 1$ that promotes out of phase oscillations of Wnt and Notch. The anterior side has a bistable behavior that forms somites with $k_1 = 4$. In the middle part of the tail, intermediate values of $k_1 = 2.3$ promote in-phase oscillations of Wnt and Notch that propagate in the form of waves due to an excitable behavior, showed in the simulation by an overlapping region of high Wnt and Notch that moves anteriorly before freezing. This model is able to form more phase waves in the expanded posterior mutant and it is robust to noise.

**Movie 9: The Clock and Wavefront Self-Organizing model can capture the changes in phase wave width observed in vivo**
The CWS model can be adjusted to incorporate a graded modulation of $k_1$ instead of a step-wise change to capture the changes in phase wave width observed in vivo.

**Movie 10: Virtual dissection of the tailbud in the CWS model**
Movie showing the effect of tailbud removal in a simulation using the Clock and Wavefront Self-Organizing model. Following the separation of the posterior region, the tailbud continues to grow and to generate new phase waves that travel anteriorly. In the anterior dissected part, preexisting phase waves persist and propagate anteriorly, driven by the excitable behavior in the intermediate region (light gray section), until they stop when entering the bistable region (dark gray section). These findings align with experimental observations reported in [23]. Importantly, in the anterior dissected part, the propagation of straight waves without rotational movements occurs independently of the homogeneous oscillations at the posterior tailbud tip.

**Movie 11: Middle explant of the The Clock and Wavefront Self-Organizing model recapitulates experimental findings**
Simulated explant obtained from the middle part of the tail in the Clock and Wavefront Self-Organizing model. The simulation recapitulates the circular Notch signaling waves that propagate from the center of the colony towards the periphery observed in experiments [10]. Simulated Wnt ($u$) and Notch ($v$) signaling are shown side-by-side. The virtual explant is generated by making a radial projection of average values along the y-axis for the middle part of the tail, with cells having $k_1 = 2.3$ that generate circular wave patterns due to two spiral centers that rotate in opposite directions.

This type of pattern depends on initial conditions and arise in 37 % (11/30) of simulations (Figure S17).

**Movie 12: Explant of the whole posterior tail in the The Clock and Wavefront Self-Organizing model recapitulates experiments**
Simulated explant obtained from the whole posterior part of the tail in the Clock and Wavefront Self-Organizing model. Simulated Wnt ($u$) and Notch ($v$) signaling are shown side-by-side. The virtual explant is generated by making a radial projection of average values along the y-axis for the whole posterior part of the tail, with cells in the center having $k_1 = 1.0$ and outermost cells having $k_1 = 2.3$. In agreement with experiment [7], oscillations at the center of the explant propagate towards the periphery.

**Movie 13: Counterclockwise waves in mixed cell explants**
Mixed explants from the CWS and CG model with diffusion. Simulated $u$/Wnt and $v$/Notch signaling are shown side-by-side. Mixed explants of the Clock and Wavefront Self-Organizing (CWS) model generate periodic phase waves with a sustained rotating pattern due to the excitable behavior of the cells. The Clock and Gradient (CG) model allows local synchronizations between cells to promote rotating patterns.

#### References


[1] Alexander Aulehla, Christian Wehrle, Beate Brand-Saberi, Rolf Kemler, Achim Gossler, Benoit Kanzler, and Bernhard G. Herrmann. Wnt3a plays a major role in the segmentation clock controlling somitogenesis. *Developmental Cell*, 4:395–406, 2003.

[2] Ahmet Ay, Stephan Knierer, Adriana Sperlea, Jack Holland, and Ertuğrul M. Özbudak. Short-lived her proteins drive robust synchronized oscillations in the zebrafish segmentation clock. *Development*, 140:3244–3253, 8 2013.

[3] James Cotterell, Alexandre Robert-Moreno, and James Sharpe. A local, self-organizing reaction-diffusion model can explain somite patterning in embryos. *Cell Systems*, 1:257–269, 2015.

[4] M. C. Cross and P. C. Hohenberg. Pattern formation outside of equilibrium. *Reviews of Modern Physics*, 65:851, 1993.

[5] Richard J. Field and Richard M. Noyes. Oscillations in chemical systems. iv. limit cycle behavior in a model of a real chemical reaction. *The Journal of Chemical Physics*, 60:1877–1884, 1974.

[6] Richard FitzHugh. Impulses and physiological states in theoretical models of nerve membrane. *Biophysical Journal*, 1:445–466, 1961.

[7] Alexis Hubaud, Ido Regev, L. Mahadevan, and Olivier Pourquié. Excitable dynamics and yap-dependent mechanical cues drive the segmentation clock. *Cell*, 171:668–682.e11, 2017.

[8] Anna Kicheva, Periklis Pantazis, Tobias Bollenbach, Yannis Kalaidzidis, Thomas Bitting, Thomas Jülicher, and Marcos González-Gaitán. Kinetics of morphogen gradient formation. *Science*, 315, 1 2007.

[9] Chandrashekar Kuyyamudi, Shakti N. Menon, and Sitabhra Sinha. Morphogen-regulated contact-mediated signaling between cells can drive the transitions underlying body segmentation in vertebrates. *Physical Biology*, 19:16001, 2022.

[10] Volker M. Lauschke, Charisios D. Tsiairis, Paul François, and Alexander Aulehla. Scaling of embryonic patterning based on phase-gradient encoding. *Nature*, 493:101–105, 2013.

[11] Julian Lewis. Autoinhibition with transcriptional delay: A simple mechanism for the zebrafish somitogenesis oscillator. *Current Biology*, 13:1398–1408, 2003.

[12] Mitsuhiro Matsuda, Hanako Hayashi, Jordi Garcia-Ojalvo, Kumiko Yoshioka-Kobayashi, Ryoichiro Kageyama, Yoshihiro Yamanaka, Makoto Ikeya, Junya Toguchida, Cantas Alev, and Miki Ebisuya. Species-specific segmentation clock periods are due to differential biochemical reaction speeds. *Science*, 369:1450–1455, 2020.

[13] A. Pillarisetti, H. Ladjal, A. Ferreira, C. Keefer, and J.P. Desai. Mechanical characterization of mouse embryonic stem cells. pages 1176–1179. IEEE, 9 2009.

[14] I. Prigogine and R. Lefever. Symmetry breaking instabilities in dissipative systems. ii. *The Journal of Chemical Physics*, 48:1695–1700, 1968.

[15] I. Prigone. Time, structure and fluctuations. *Science*, 201:777–785, 1978.

[16] Rajeev Singh and Sitabhra Sinha. Spatiotemporal order, disorder, and propagating defects in homogeneous system of relaxation oscillators. *Physical Review E - Statistical, Nonlinear, and Soft Matter Physics*, 87:012907, 2013.

[17] Katharina F. Sonnen, Volker M. Lauschke, Julia Uraji, Henning J. Falk, Yvonne Petersen, Maja C. Funk, Mathias Beaupeux, Paul François, Christoph A. Merten, and Alexander Aulehla. Modulation of phase shift between wnt and notch signaling oscillations controls mesoderm segmentation. *Cell*, 172:1079–1090.e12, 2018.

[18] Yoshiki Takashima, Toshiyuki Ohtsuka, Aitor González, Hitoshi Miyachi, and Ryoichiro Kageyama. Intronic delay is essential for oscillatory expression in the segmentation clock. *PNAS*, 108:3300–3305, 2 2011.

[19] P. P. L. Tam. The control of somitogenesis in mouse embryos. *Development*, 65:103–128, 10 1981.

[20] Reza Torabi and Jörn Davidsen. Pattern formation in reaction-diffusion systems in the presence of non-markovian diffusion. *Phys. Rev. E*, 100:052217, 2019.

[21] Charisios D. Tsiairis and Alexander Aulehla. Self-organization of embryonic genetic oscillators into spatiotemporal wave patterns. *Cell*, 164:656–667, 2016. Somitogenesis.

[22] A. M. Zhabotinsky and A. N. Zaikin. Autowave processes in a distributed chemical system. *Journal of Theoretical Biology*, 40:45–61, 1973.

[23] Ece Özelçi, Erik Mailand, Matthias Rüegg, Andrew C. Oates, and Mahmut Selman Sakar. Deconstructing body axis morphogenesis in zebrafish embryos using robot-assisted tissue micromanipulation. *Nature Communications*, 13, 12 2022.
